# A time-series analysis of blood-based biomarkers within a 25-year longitudinal dolphin cohort

**DOI:** 10.1101/2022.06.28.497095

**Authors:** Aaditya V. Rangan, Caroline C. McGrouther, Nivedita Bhadra, Stephanie Venn-Watson, Eric D. Jensen, Nicholas J. Schork

## Abstract

1

Causal interactions and correlations between clinically-relevant biomarkers are important to understand, both for informing potential medical interventions as well as predicting the likely health trajectory of any individual as they age. These interactions and correlations can be hard to establish in humans, due to the difficulties of routine sampling and controlling for individual differences (e.g., diet, socio-economic status, medication). Because bottlenose dolphins are long-lived mammals that exhibit several age-related phenomena similar to humans, we analyzed data from a well controlled 25-year longitudinal cohort of 144 dolphins. The data from this study has been reported on earlier, and consists of 44 clinically relevant biomarkers. This time-series data exhibits three starkly different influences: (A) directed interactions between biomarkers, (B) sources of biological variation that can either correlate or decorrelate different biomarkers, and (C) random observation-noise which combines measurement error and very rapid fluctuations in the dolphin’s biomarkers. Importantly, the sources of biological variation (type-B) are large in magnitude, often comparable to the observation errors (type-C) and larger than the effect of the directed interactions (type-A). Attempting to recover the type-A interactions without accounting for the type-B and type-C variation can result in an abundance of false-positives and false-negatives. Using a generalized regression which fits the longitudinal data with a linear model accounting for all three influences, we demonstrate that the dolphins exhibit many significant directed interactions (type-A), as well as strong correlated variation (type-B), between several pairs of biomarkers. Moreover, many of these interactions are associated with advanced age, suggesting that these interactions can be monitored and/or targeted to predict and potentially affect aging.

**Author Summary:** The body is a very complicated system with many interacting components, the vast majority of which are practically impossible to measure. Furthermore, it is still not understood how many of the components that we *can* measure influence one another as the body ages. In this study we try and take a small step towards answering this question. We use longitudinal data from a carefully controlled cohort of dolphins to help us build a simple model of aging. While the longitudinal data we use does measure many important biomarkers, there are obviously a much larger number of biomarkers that haven’t been measured. Our simple model accounts for these ‘missing’ measurements by assuming that their accumulated effect is similar to a kind of ‘noise’ often used in the study of complicated dynamical systems. With this simple model we are able to find evidence of several significant interactions between these biomarkers. The interactions we find may also play a role in the aging of other long-lived mammals, and may be worth investigating further to better understand human aging.

## 3 Introduction

A better understanding of the biology of aging can help us discover strategies to stay healthier longer [1, 2]. Studying aging directly in humans has many advantages, and can help pinpoint some of the mechanisms responsible for predicting and controlling aging rates [3, 4, 5, 6, 7]. Unfortunately, human studies have several disadvantages, including difficulties acquiring regular samples, as well as controlling for the differences between individuals (e.g., diet, socioeconomic status and medication) [6, 8, 9]. Many of these limitations can be overcome by focusing on short-lived animals such as worms, flies and mice [10, 11], but it is important to complement this work with studies of animals that have lifespans similar to humans (i.e., with evolutionary adaptations that allow them to live 50 or more years long).

Bottlenose dolphins (*Tursiops truncatus*) provide a useful model organism, as they are long-lived mammals which perform complex social and cognitive tasks over the course of their lives [12, 13, 14, 15]. Importantly, dolphins also share many genetic and biogerontological qualities with humans [16, 14, 17, 18, 19, 20, 21, 22]. For example, dolphins exhibit some similarities to humans regarding cellular function [23], and reduce their energy expenditure as they age [24]. Dolphins also develop several age-related conditions – such as chronic inflammation and insulin-resistance – with clinical signs similar to humans [18, 22, 25, 26, 27], while also developing histological lesions similar to certain human disorders [26, 27, 28, 29, 30].

In this manuscript we analyze data taken from a longitudinal cohort of 144 US Navy bottlenose-dolphins. These dolphins were cared for over three generations and were routinely sampled to asess the trajectory of 44 clinically relevant biomarkers over their lifespan, as described in [20, 31, 32]. The diet and environment of these dolphins was carefully controlled, resulting in the dolphins living an average of 32.5 years (c.f. wild dolphins live an average of 20 years) [33, 34]. The uniformity of environment, diet and health-care within this this cohort implies that age-related differences between dolphins might be due to inherent or genetic factors influencing senescence [35].

This data-set was used previously to demonstrate differences in aging-rates between individual dolphins [35]. Motivated by these results, we further analyze the trajectories of each of the biomarkers for each of the dolphins within this data-set, searching for hints of any causal interactions between the measured biomarkers. In this analysis it is crucial to account for the shared fluctuations (termed ‘shared variation’) between biomarkers, as these shared fluctuations typically dwarf the more subtle causal interactions we are trying to find. To account for this shared variation, we model the longitudinal data as a linear stochastic-differential-equation (SDE), extracting model parameters as indicators of which biomarkers might affect one another. After doing so, we are able to identify interactions that are associated with aging. These interactions may play a role in the biology of aging, and are prime candidates for further investigation.

## 4 Results

### 4.1 Motivation for the Model

Obviously, the body is a very complicated dynamical system [36]. Any set of biomarkers can only ever tell part of the story, and there will always be an abundance of unmeasured factors that play an important role in the trajectory of all the measured biomarkers. Given any particular subset of (observed) biomarkers, we typically expect the changes in those biomarkers to come from three different categories.

**type-A: Directed interactions.** Any particular biomarker may play a direct role in influencing the dynamics of other biomarkers. For example, one biomarker might ‘excite’ or ‘inhibit’ another, with an increase in the first promoting an increase (or, respectively, decrease) in the second.
**type-B: Shared biological variation.** As the system evolves, we expect a large number of unmeasured biomarkers to affect each of the measured biomarkers. Generally, this multitude of unmeasured effects will accumulate, giving rise to a combined effect that is essentially unpredictable given the measurements at hand. Collectively, the influence of these unmeasured effects can seem random.
**type-C: Observation-noise.** To complicate things further, our measurements of each observed biomarker may not be perfectly accurate. Typically there are additional sources of ‘observation-noise’ which introduce random variation to our measurements. This type-C variation can include both measurement errors as well as true biological fluctuations that are much more rapid than the typical time-interval of the experimental measurements themselves (e.g,. hourly variations based on the individual’s activity).

To understand how large a role each category plays, we can look at the changes in each of the biomarkers within the dolphin data-set. One example for the biomarker ‘Alkaline Phosphatase’ (AlkPhos) is shown in Fig 1. This figure illustrates the distribution of measured variable-increments (between one time-point and the next) for this biomarker. The large variation in these increments indicates that the type-B and type-C effects play a large role. Furthermore, there is a striking linear relationship between (i) the magnitude of the measured time-increment and (ii) the variance in the measured variable-increments. This linear relationship indicates that a significant component of the type-B variation is similar to the Brownian processes commonly seen in high-dimensional dynamical systems [37]. Such a Brownian process – also called a ‘stochastic drive’ – can provide a source of shared variation which correlates different biomarkers. Finally, we remark that the variation in variable-increments is still significant even when the measured time-increment is 0. In other words, multiple repeated measurements of the same dolphin may not always return the same value. This indicates that there is a significant source of type-C variation in our measurements.

**Figure 1:**
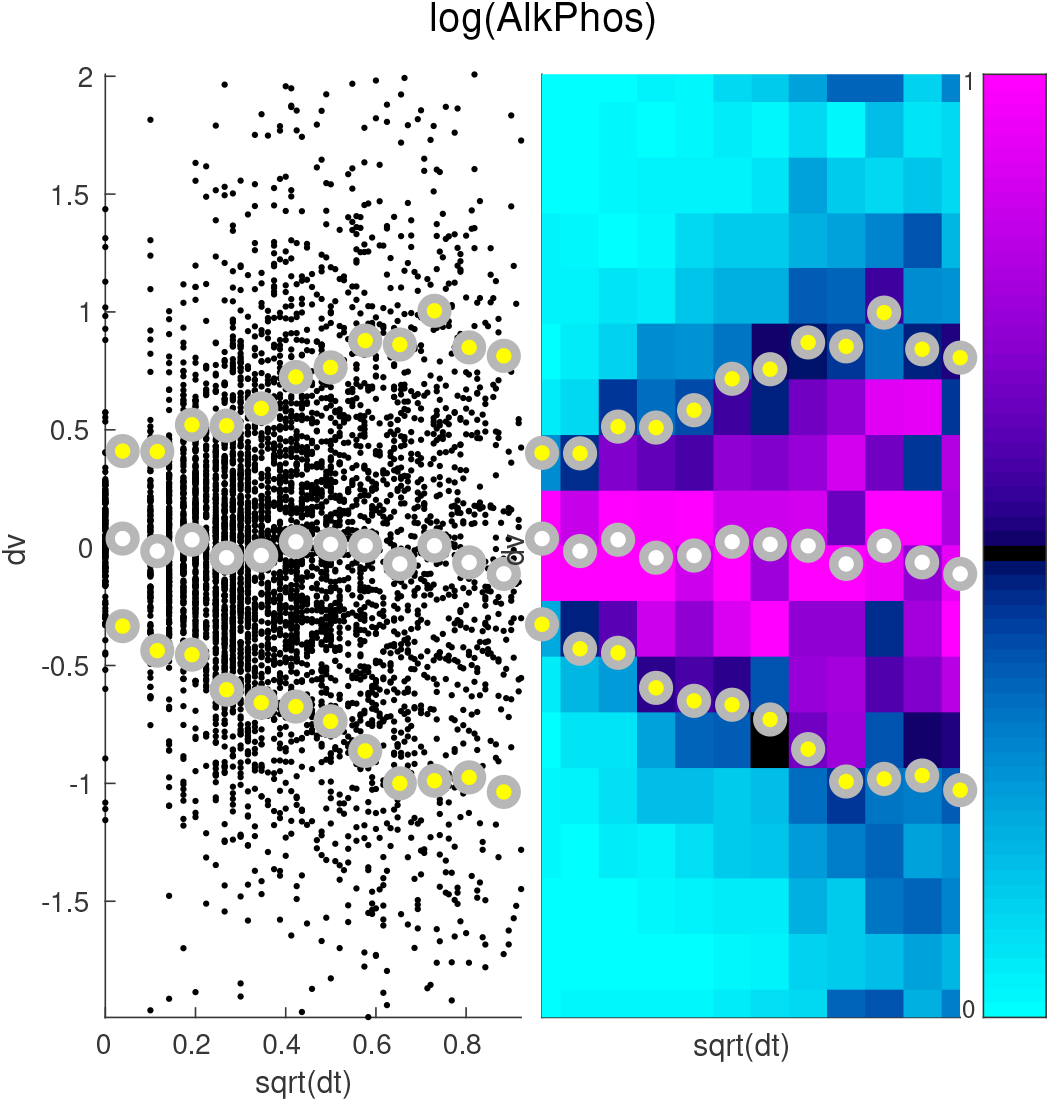
Here we illustrate the distribution of increments associated with Alkaline Phosphatase (AlkPhos). On the left we show a scatterplot of these increments. Each small black point in the background represents a pair of time-adjacent measurements in a single dolphin, with the square-root of the time-increment (in years) along the horizontal, and the change in AlkPhos along the vertical. The time-increments are then divided into 12 bins, and the mean (white dot) plus and minus one standard-deviation (yellow dots) are displayed in the foreground. On the right we show the same data, with the 12 time-bins shown as vertical strips. Each vertical strip is further divided into 18 boxes, which are colored by the number of increments within that time-bin which fall into that box (i.e., a histogram for each time-bin, scaled to have maximum 1, colored as shown in the colorbar to the far right). Note that the distribution of increments for each time-bin becomes wider as the time-increment increases. Importantly, the standard-deviation increases roughly linearly with the square-root of the time-increment (i.e., the variance increases roughly linearly with the time-increment), as expected from a Brownian-process. Note also that the variance is nonzero even when the time-increment is 0, implying that there must be an extra source of variation in addition to this Brownian-process.

The qualitative aspects of Fig 1 are typical for this dataset; Figs 2 and 3 illustrate the distribution of variable-increments for two other measured biomarkers. Note that, once again, we see that the variation in variable-increments is large, and that there is evidence for both type-B and type-C variation. For some biomarkers we expect the type-B variation to be larger than the type-C variation, whereas for other biomarkers we expect the type-C variation to dominate (e.g., compare Figs 2 and 3).

**Figure 2:**
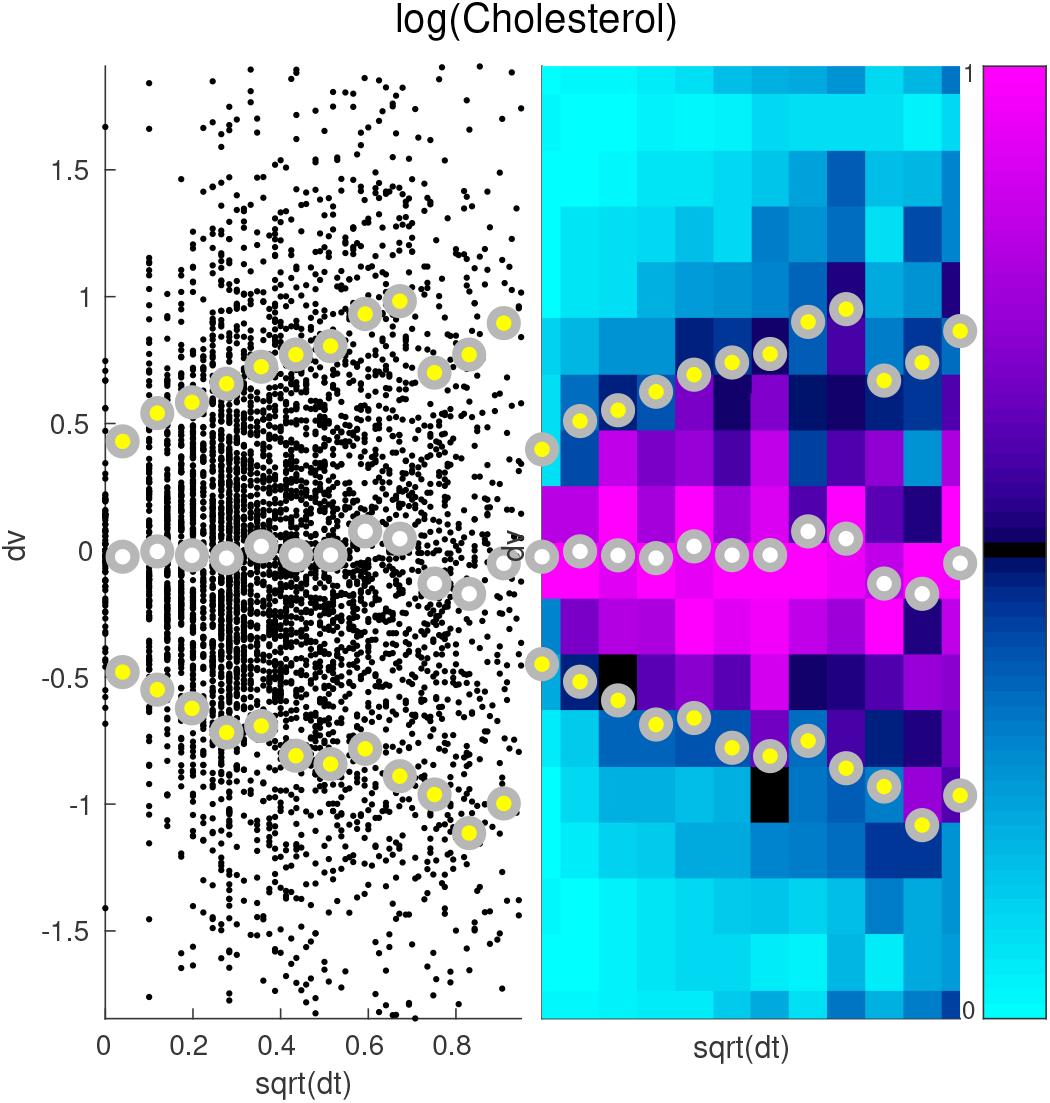
Here we illustrate the distribution of increments associated with Cholesterol. The format for this figure is the same as Fig 1.

**Figure 3:**
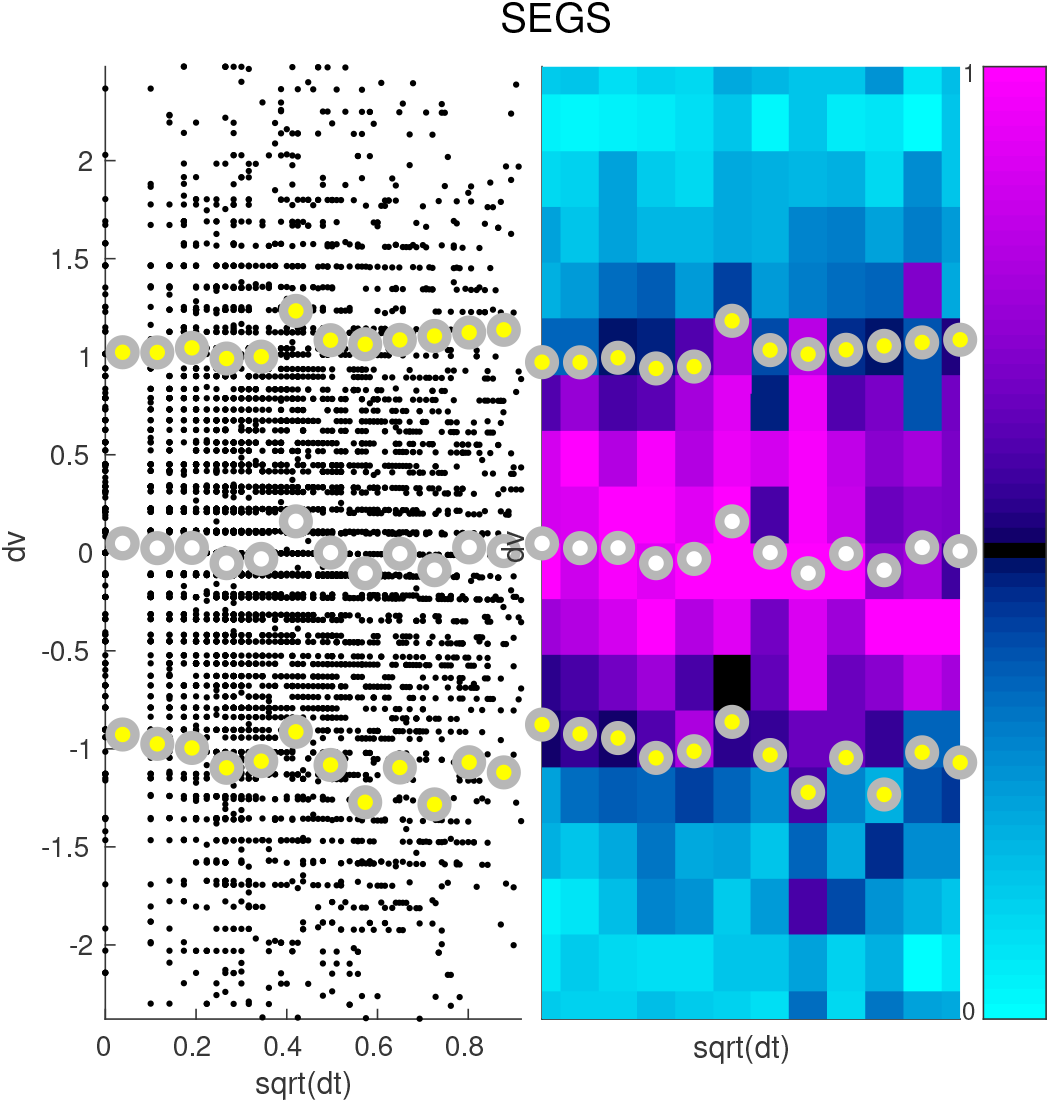
Here we illustrate the distribution of increments associated with Segmented Neutrophils (SEGS). The format for this figure is the same as Fig 1.

In the next section we’ll introduce our model for the dynamics. Within this model we’ll model the type-A directed interactions as simple linear interactions (see (1) below). Motivated by the observations above, our model will also include terms to account for type-B and type-C variations. we’ll model the type-B variations as an (anisotropic) Brownian-process (see (1) below), and we’ll model the observation-noise as an (anisotropic) Gaussian with time-independent variance [38] (see (2) below).

### 4.2 Model Structure

Given the motivation above, we can model the evolution of any *d* specific variables over the time-interval [*t*, *t*′] using a simple linear stochastic-differential-equation (SDE) of the following form:

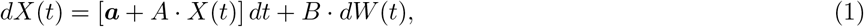

where 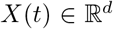 represents the *d*-dimensional vector-valued solution-trajectory at the initial time *t*, the time-increment *dt* = *t*′ – *t* represents the difference between the initial time *t* and the final time *t*′, and 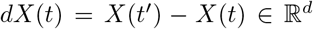 represents the vector of variable-increments between times *t* and *t*′. We further assume that we do not measure *X*(*t*) directly, but rather some *Y*(*t*) which depends on *X*(*t*) and which also incorporates the type-C observation-noise:

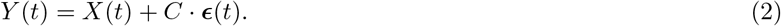

In (1) and (2) there are several terms on the right hand side which contribute to the observed dynamics.

**Baseline velocity:** The vector 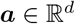 corresponds to a constant ‘velocity’ for each of the variables. This represents the rates at which each variable would increase, were it not for the other influences within the system.
**Directed interactions:** The matrix 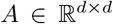 represents the type-A interactions between variables. These directed interactions are modeled as linear effects; the value of a ‘source’ variable *v*′ will influence the rate at which the ‘target’ variable *v* increases at time *t* via the factor 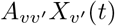. Each of these type-A interactions can be thought of as part of a pathway or larger network describing how the variables influence one another. These directed interactions are often called ‘deterministic interactions’, as they do not explicitly involve any randomness.
**Stochastic drive:** The matrix *B* represents the biological variation (due to an accumulation of independent unmeasured effects) expected to affect the model variables as they evolve. The source of the type-B variation is modeled by the noise *dW*(*t*) (i.e., Brownian-increments) which are each drawn independently from the Gaussian distribution 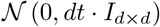. The matrix 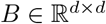 controls the correlations between the noise received by the different variables [37]. Put another way, *dtBB*^⊤^ is the covariance matrix of biological variation for this model, analogous to the variance-covariance matrix in quantitative genetics [39]. This biological variation is often called a ‘stochastic drive’ to disambiguate it from the type-C observation-noise mentioned below.
**Observation-noise:** The matrix *C* in (2) represents both the noise inherent to the observations, as well as the effects of very rapid unpredictable fluctuations in the biomarkers. Each vector 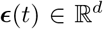 at each observation-time is drawn (independently) from the Gaussian distribution 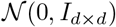. The matrix 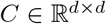 controls the level of correlations within the type-C errors; the covariance of this observation-error is given by the symmetric-matrix *CC*^⊤^.

Note that the observation-times are not necessarily unique: *t*′ could very well equal *t*, and *dt* may equal 0. In this situation *dX*(*t*) ≡ **0**, and so *X*(*t*′) will equal *X*(*t*). However, because of the type-C errors, *Y*(*t*′) will in general be different from *Y*(*t*).

While we could, in principle, fit an SDE to all the variables simultaneously (with a large *d*), we opt to consider only pairwise combinations of variables (with *d* = 2 in each case). Mathematically, each such pairwise fit (for variables *v* and *v*′) corresponds to approximating the system’s trajectory after projecting the full system onto the subspace spanned by those two variables. We make this decision for three reasons.

**Increase Power:** First and foremost, the number of model parameters, as well as the number of observations required to accurately estimate these parameters, both increase as *d* increases. We will have the most statistical power when *d* is low. This topic is discussed in more detail within the Supplementary section 3.3.
**Avoid Redundancy:** Second, redundancies in any subset of variables (e.g., GFR and Creatinine, or MCH and MCV) can easily create the illusion of large interactions between the redundant set and other variables. These spurious interactions will be statistically, but not biologically significant. In mathematical terms, these redundancies will result in a poorly conditioned inverse problem, which we want to avoid [40].
**Easily Understand:** By considering only pairwise interactions, the results can be interpreted more easily. The interactions between any pair of variables can then be considered (and used to make predictions) even in situations where the other variables aren’t accessible, or haven’t been measured.

We acknowledge that there are many strategies for modeling the interactions in this longitudinal data-set (see, e.g., [41, 42, 43, 44, 45]). For example, there are many similarities between our model and vector-autogregression [46]. Indeed, if the time-intervals *dt* in this data-set were all of equal size, then our method would be equivalent to one-step vector-autogregression; the only structural difference being that in such a scenario the type-B and type-C variation would be indistinguishable from one another. We opt for the SDE proposed above because (i) it has relatively few parameters, (ii) it is easy to understand, and (iii) it can easily accommodate the irregular time-intervals and occasional missing elements within this data-set.

### 4.3 Model Interpretation

We use a generalized regression (see Supplementary section 3) to fit the longitudinal data for each pair of biomarkers *v* and *v*′ with the simple dynamical system (SDE) above. After fitting the model parameters we interpret *A* and *BB*^⊤^ as having potential biological significance.

The entries *A_vv′_* indicate the directed interactions (i.e., the deterministic effects) from biomarker *v*′ to biomarker *v*. If *A_vv′_* is positive, then we expect an increase in biomarker *v*′ to typically precede a subsequent increase in biomarker v; we conclude that *v*′ ‘excites’ *v*. Conversely, if *A_vv′_* is negative, then we expect that *v*′ will ‘inhibit’ *v*; an increase in *v*′ will typically precede a subsequent decrease in *v*.

The symmetric entries [*BB*^⊤^]_*vv*′_ = [*BB*^⊤^]_*v′v*_ indicate the shared biological variation (i.e., the stochastic drive) expected to influence both biomarkers *v* and *v*′. If [*BB*^⊤^]_*vv*′_ is positive, then *v* and *v*′ will experience correlated stochastic input which will give rise to correlated (but not causal) fluctuations between these two biomarkers. Conversely, if [*BB*^⊤^]_*vv*′_ is negative, then *v* and *v*′ will experience anti-correlated stochastic input which will produce anti-correlated (but again, not causally-linked) fluctuations between the two biomarkers. Because we fit the SDE to each pair of biomarkers individually, the correlations (or anti-correlations) within this stochastic drive to the pair *v*, *v*′ can result from the accumulated effects of the other measured biomarkers (other than *v*, *v*′), as well as from biomarkers which weren’t included in the study at all.

To illustrate these kinds of relationships, we fit the SDE in (1),(2) to the two biomarkers ‘Iron’ and ‘Mean corpuscular hemoglobin’ (MCH) across all observation-times across all dolphins. Referring to Iron and MCH as *v*′ and *v*, respectively, we find that *A_vv′_* is significantly positive and *A_v′v_* is significantly negative (with uncorrected *p*-values *p*_0_ < 1e – 11 and *p*_0_ < 1e – 4 respectively, see Methods). Moroever, we find that [*BB*^⊤^]_*vv*′_ is significantly positive (*p*_0_ < 1e – 50). As we’ll discuss below, the type-B correlations are quite a bit larger (in magnitude) than the type-A interactions for these two biomarkers (note the difference in *p*-values), and an accurate estimate of *A*_*vv*′_ can only be obtained after estimating [*BB*^⊤^]_*vv*′_.

The directed interactions *A* can be visualized using the phase-diagram of the linear differential-equation associated with *A* (i.e., by ignoring the effect of *B* in (1)). This phase-diagram is shown on the left of Fig 4, with the directed interactions giving rise to the flow vectors in the background (grey). For this differential-equation a surplus of Iron will drive an increase in MCH, while a surplus of MCH will drive a decrease in Iron. This combination of excitation and inhibition will tend to give rise to ‘counterclockwise’ motion within the Iron-MCH phase-plane (indicated by the directions of the grey arrowheads).

**Figure 4:**
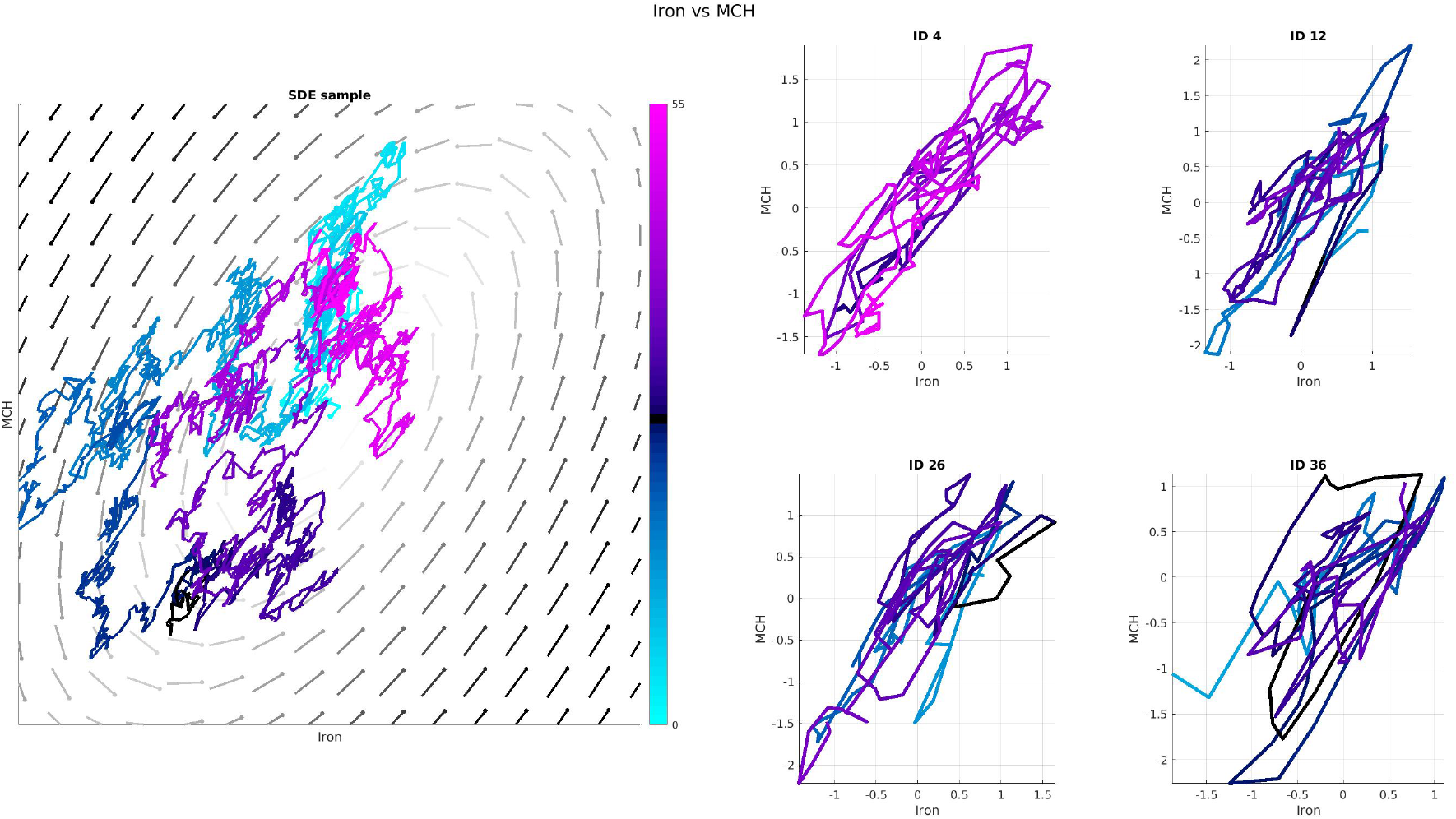
Here we illustrate the directed interactions between Iron (horizontal) and mean corpuscular hemoglobin (MCH) (vertical). The left-hand-side shows the phase-diagram associated with *A*, as well as a sample-trajectory from the SDE (including *B*). Other sample trajectories are shown in Supplementary Fig 10. The right-hand-side shows several subplots from various dolphins (after kalman filtering to reduce the observation error). The colorscale refers to age (see inset colorbar).

The subplot on the left of Fig 4 also shows a finely sampled trajectory *X*(*t*) of the SDE (which includes the effect of the nonzero *B*). This trajectory is colored by age, from 0yr (cyan) to 55yr (magenta) (see adjacent colorbar). Note that the sample-trajectory often deviates from the counterclockwise course suggested by the directed interactions, often meandering around due to the stochastic drive provided by *B*. The sample trajectory shown here is just one possible trajectory for the SDE; other sample trajectories are shown in Supplementary Fig 10. Subplots illustrating the (kalman filtered) trajectories of several individual dolphins are shown on the right of this figure, using the same colorscale. Similar to the sample trajectory from the SDE, these experimental trajectories also meander around, while exhibiting a global tendency to prefer counterclockwise loops over clockwise ones.

The type-B and type-C variations in these two biomarkers are shown in Fig 5. To illustrate these sources of variation, we take all the time-increments *dt* used to fit the SDE (across all pairs of adjacent observation-times and all dolphins), and divide them into 10 percentile-bands. For each percentile-band of *dt* we construct a scatterplot of the observed variableincrements 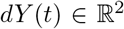 for these two biomarkers. We fit the distribution of variable-increments in each percentile-band with a Gaussian, and indicate the 1- and 2-standard-deviation contours of this Gaussian with bright- and pale-yellow ellipses, respectively. The covariance-matrix *CC*^⊤^ is approximately equal to the covariance of the scatterplot shown in the upper-left, with *dt* ~ 0. The covariance-matrix for each of the other subplots is roughly given by *dtBB*^⊤^ + *CC*^⊤^; as *dt* increases the variable-increments become more and more correlated, indicating that [*BB*^⊤^]_*vv*′_ is positive (i.e., these two biomarkers are influenced by a shared and correlated stochastic drive).

**Figure 5:**
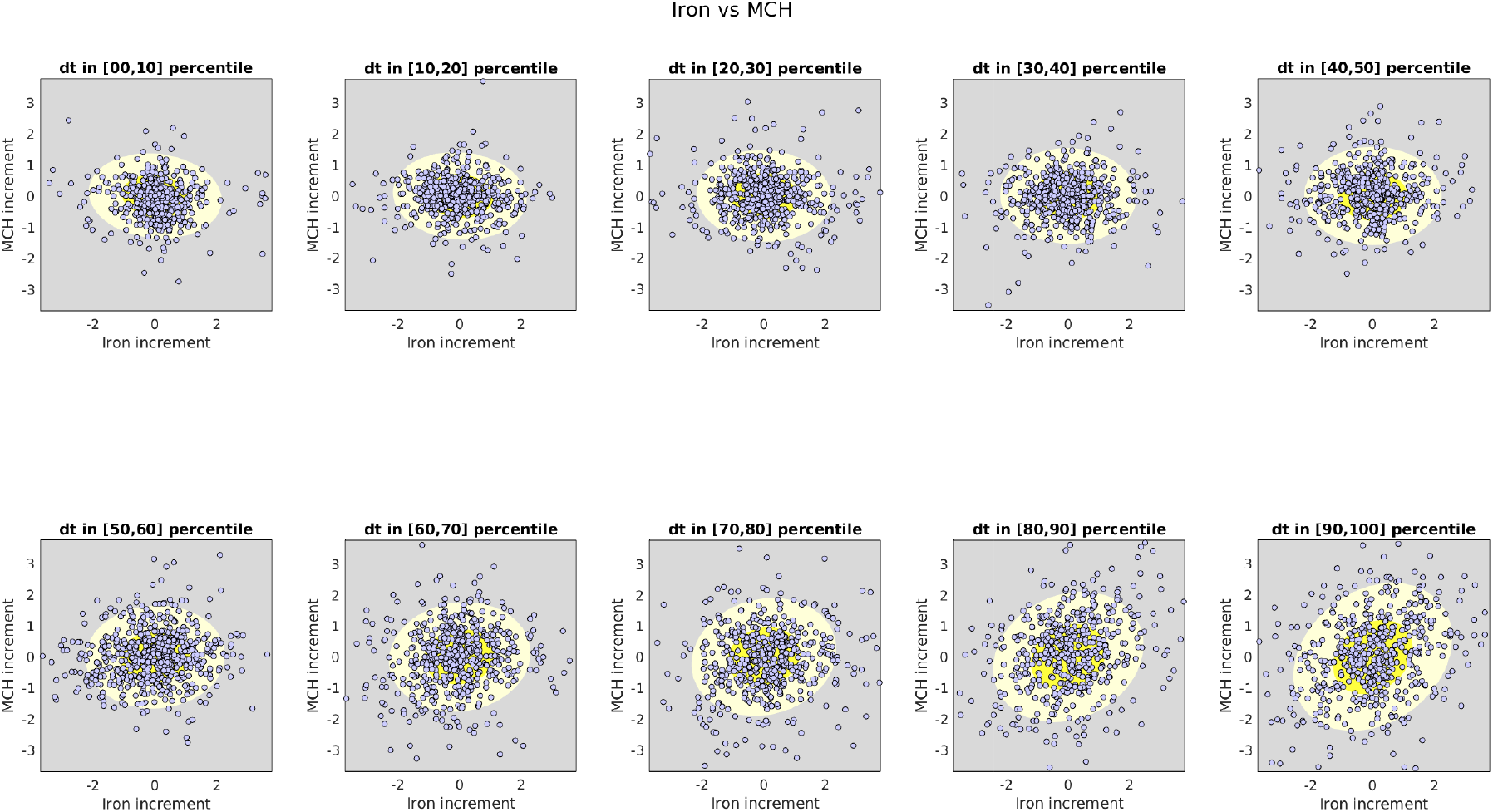
Here we illustrate the type-B and type-C variations associated with Iron (horizontal) and MCH (vertical). Each subplot shows a scatterplot of variable-increments for a percentile-band of observed time-increments. The yellow ellipses correspond to standard-deviation contours of the best-fit Gaussian distribution within each scatterplot.

For these two biomarkers the type-B and type-C variations contribute heavily, dominating the evolution of the trajectory of the SDE. Specifically, the term *B* in (1) is several times larger than the matrix *A*. The term *C* in (2) is also large, contributing observation-noise which is a few times larger (on average) than the type-A interactions. This is quite common, and most biomarker-pairs exhibit a similar phenomenon.

If we were to ignore the type-B and/or type-C interactions when fitting this data (i.e., setting *B* ≡ 0 in (1) or *C* ≡ 0 in (2)), then the correlations in these variations would impact our recovery of *A*, giving rise to inaccurate estimates of *A*_*vv*′_. These spurious estimates for *A*_*vv*′_ would, in turn, give rise to an abundance of false-positives and false-negatives when estimating the significance of the directed interactions across the biomarker-pairs within the data-set. By accounting for these type-B and type-C terms within the SDE, we believe that we can accurately estimate the size and direction of the type-A interactions.

To summarize, we expect Iron and MCH to exhibit strong correlated (but not necessarily causal) fluctuations due to shared input coming from the accumulated effect of other (potentially unmeasured) biomarkers. By taking into account these strong type-B correlations, we can attempt to tease out more subtle directed type-A interactions. For these two biomarkers we do indeed find evidence of a statistically significant directed link: an increase in Iron will tend to produce a subsequent increase in MCH, while an increase in MCH will tend to produce a subsequent reduction in Iron.

### 4.4 Longitudinal Analysis

By fitting the longitudinal data with the SDE (1),(2) above, we observe multiple significant (type-A) directed interactions, as well as (type-B) shared biological variation. An illustration of the type-A interactions observed across all dolphins and ages is shown in Fig 6, while the corresponding type-B correlations are shown in Fig 7.

**Figure 6:**
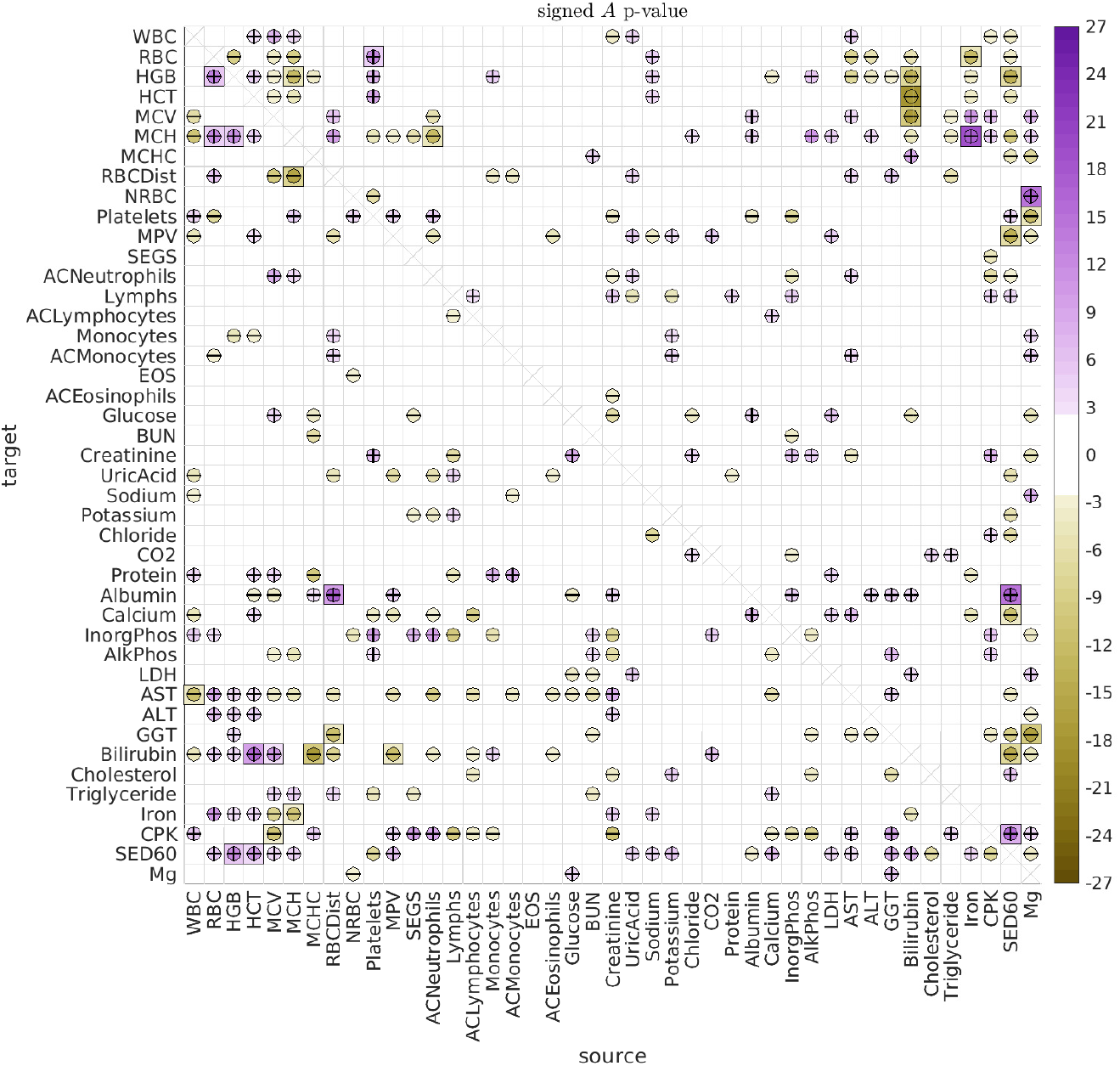
Here we illustrate the significant directed interactions *A_vv′_* observed when pooling all dolphins and all ages. The ‘source’ biomarker *v*′ is shown along the horizontal, with the ‘target’ *v* shown along the vertical. The color of each interaction indicates the log-*p*-value for that interaction, ascribed a sign according to the sign of the interaction. Thus, interactions are colored lavender if the interaction is positive, and goldenrod if the interaction is negative (see colorbar on the right). For ease of interpretation, interactions are also given a ‘+’ or ‘-’ symbol to indicate their sign. The bonferroni-corrected *p*-values *p_b_* are shown as squares, while the inscribed circles indicate the associated holm-bonferroni adjusted *p_h_*.

**Figure 7:**
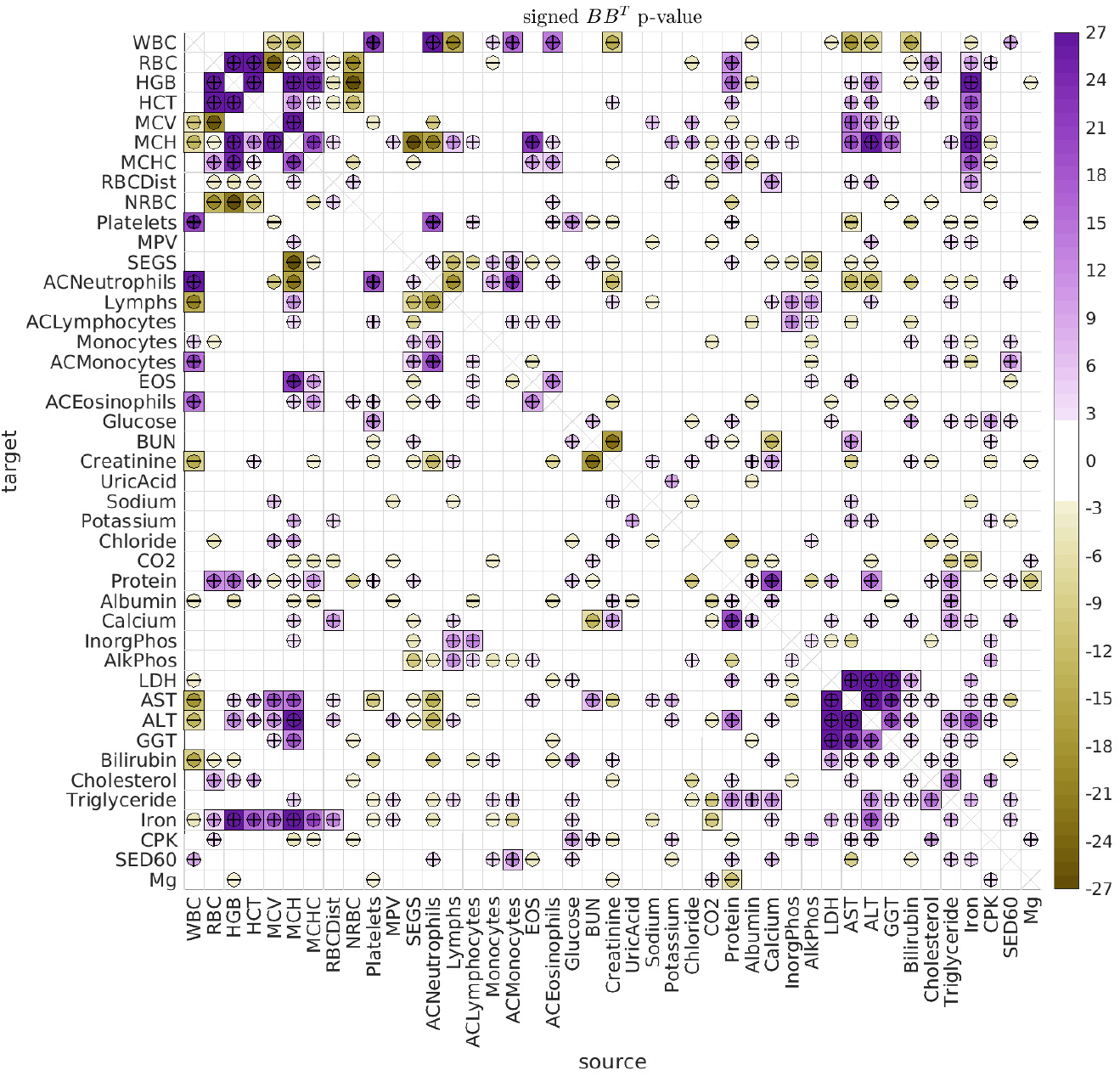
This figure has the same format as Fig 6, showing the significance of the covariance [*BB*^⊤^]_*vv*′_. As in Fig 6, these results refer to pooling all dolphins and all ages.

We first remark that the type-B correlations typically dominate the type-A interactions. We saw an example of this earlier when considering the case of Iron versus MCH, and the same general trend holds for most of the biomarker pairs. Many strong type-B correlations can be observed between the biomarkers relating to red blood cells (RBCs) and hemoglobin (HGB), such as Iron, RBC, HGB, HCT, MCV, MCH and MCHC. These correlations are to be expected, because many of these biomarkers are influenced by the same underlying biological factors [47, 48]. Similarly, we see many strong type-B correlations between the biomarkers relating to liver function, such as LDH, AST, ALT and GGT. Once again, these are all influenced by the same factors, and strong correlations are to be expected [49, 50].

Regarding the type-A interactions, we note that most of the more significant type-A interactions involve an *A_vv′_* and *A_v′v_* with opposite signs. That is, one biomarker excites the other, while being inhibited in return (see, e.g., the interaction between Iron and MCH mentioned earlier). These ‘push-pull’ pairs manifest when the observed data for biomarkers *v* and *v*′ both look like noisy versions of the same rough trajectory (e.g., meandering back-and-forth), but with one of the biomarkers ‘leading’ the other in time. The linear SDE that best fits this scenario is usually one which gives rise to stochastic oscillations with a distribution of temporal correlations fit to what was observed in the real data.

Some of the more statistically significant push-pull interactions include: (i) Iron ⇌ MCH, (ii) HGB ⇌ SED60, (iii) HCT+MCH ⇌ Bilirubin, and (iv) HGB ⇌ MCH. These interactions suggest an intricate interplay between Iron, various RBC attributes, and inflammation (for which SED60 is a proxy). These interactions are not without precedent in humans. For example, inflammation is well known to have wide-ranging effects, and can alter red blood cell clearance as well as red blood cell counts [51, 52]. The production and regulation of Heme and Hemoglobin also involves many mechanisms, and can be affected by inflammation, in turn affecting endothelial cell activation [53, 54, 55, 56, 57]. The breakdown and/or destruction of RBCs can also release damage-associated molecular patterns which can activate inflammatory pathways [58], and result in increased bilirubin levels, which in turn can trigger erythrocyte death [59].

#### Grouping the interactions

When observing the array of type-A interactions in Fig 6, it is clear that there are an abundance of ‘push-pull’ biomarker-pairs (such as Iron and MCH). Further inspection reveals that these push-pull pairs are not haphazardly scattered throughout the array (i.e., they are not ‘randomly distributed’). Instead, there are subsets of biomarkers which exhibit collective push-pull interactions.

For example, let’s define the subsets 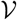 and 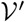 as follows. Let 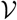 contain the biomarkers RBC, HGB, and HCT, and let 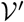 contain the biomarkers AST, MCH, Bilirubin, ALT, Sed60 and Iron. With this definition, we see that these two subsets 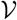 and 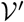 form a ‘block’ of push-pull interactions. That is, the directed interaction between any source biomarker from set 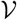 and any target biomarker from set 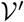 is excitatory, while the reciprocal interaction is inhibitory. These directed interactions may be related to the observed relationships between liver function and RBC elimination [60, 61, 62].

The particular block of push-pull interactions described above is quite large (comprising a set of 3 and a set of 6 biomarkers), and is substantially larger than the typical push-pull blocks one would expect if the A-type interactions were randomly reorganized (p-value < 1e – 21). There are other (less significant) push-pull blocks within the array of A-type interactions as well. To systematically search for these push-pull blocks, we use a modified version of the biclustering techniques described in [63] (see Supplementary section 4).

We can use this strategy to rearrange the biomarkers to reveal the significant push-pull blocks. As described in the Supplementary Information, many of the significant push-pull blocks involve overlapping subsets of biomarkers, and there is no single best strategy for presenting them all. One possible rearrangement, which isolates some of the larger disjoint push-pull blocks, is shown in Fig 8. In addition to the push-pull block described above (shown in the upper left of Fig 8), there are two other push-pull blocks that can be seen towards the center of Fig 8. The first is defined by 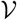 containing AlkPhos and InorgPhos, and 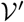 containing CPK, Platelets and BUN (p-value < 0.007). The second is defined by 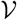 containing Creatinine and Lymphs, and 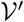 containing AlkPhos, InorgPhos, CPK and Platelets (p-value < 0.026).

**Figure 8:**
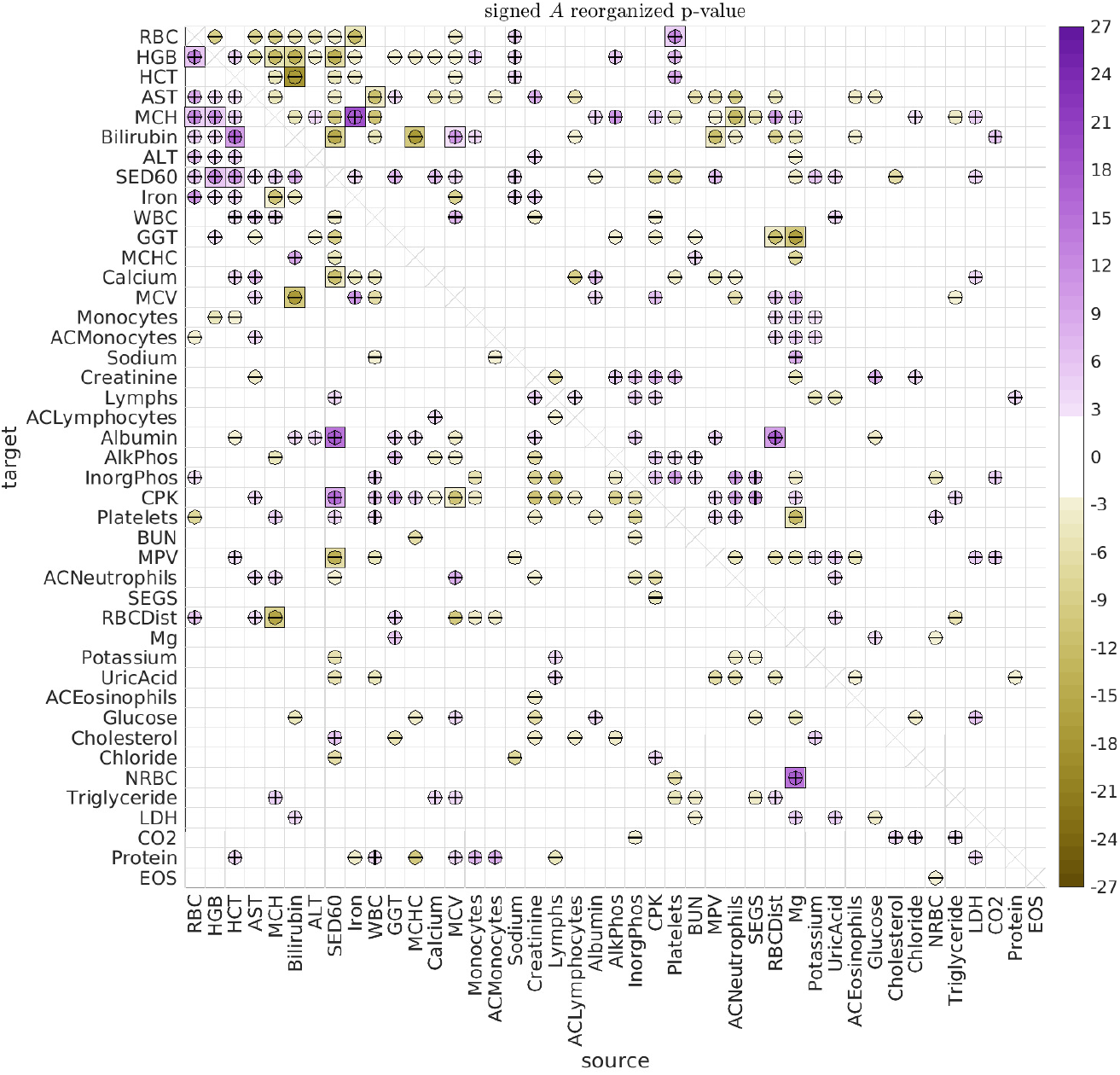
This figure has the same format as Fig 6, except that we have rearranged the biomarkers to emphasize some of the most significant distinct push-pull blocks.

When considering the type-B interactions the situation is simpler. The array of B-type interactions is symmetric, and is already quite structured. A simple spectral clustering (i.e., sorting the leading term in an eigendecomposition of *BB*^⊤^) can be used to reveal many groups of correlated biomarkers. In Fig 9 we use this strategy to rearrange the biomarkers to reveal several of the most significant (and disjoint) groups.

**Figure 9:**
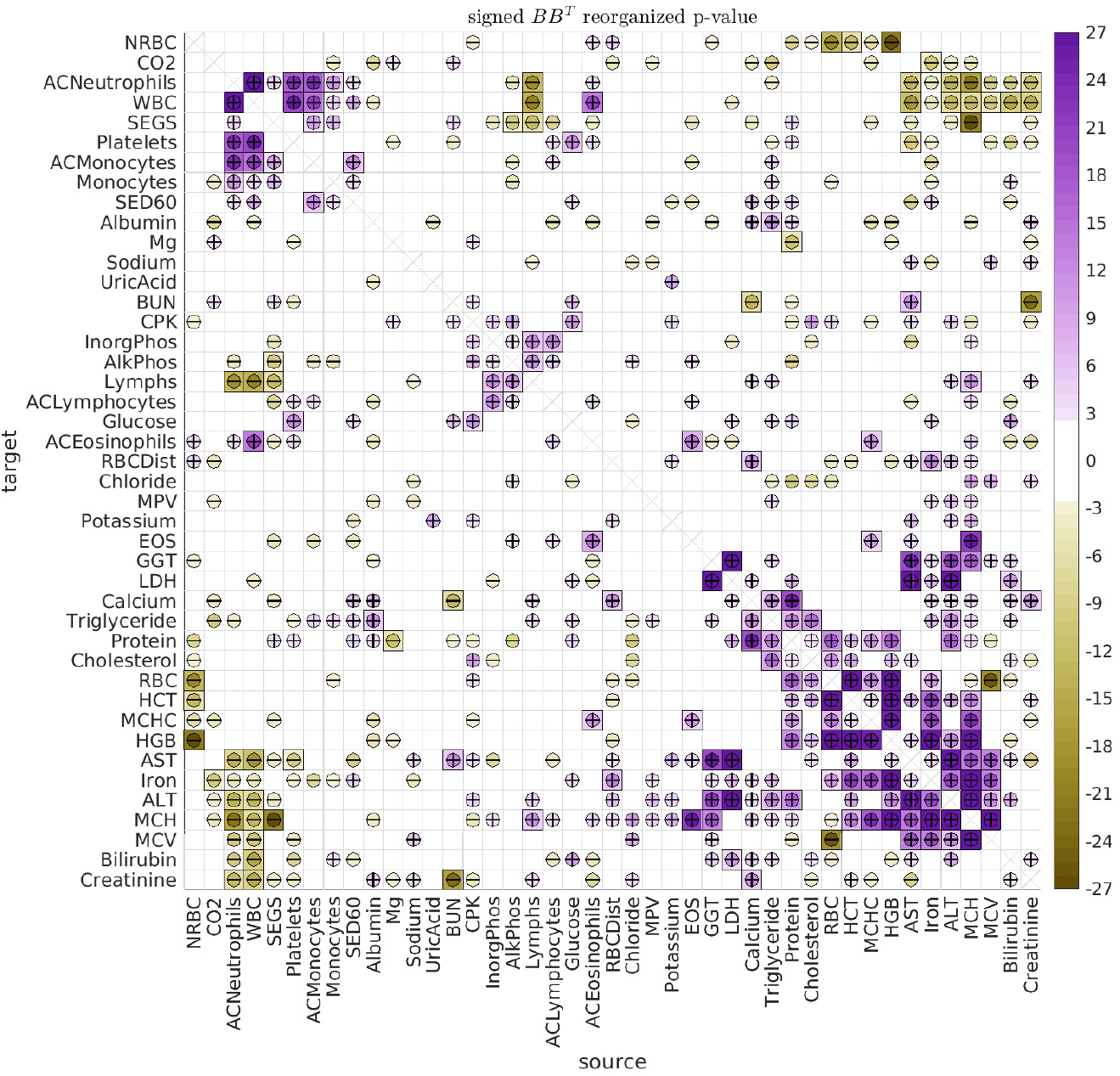
This figure has the same format as Fig 7, except that we have rearranged the biomarkers to emphasize some of the most significant clusters.

In this figure one can immediately see several different subsets of biomarkers which exhibit high levels of intra-group correlation. For example, the group of biomarkers in the bottom right corner of Fig 9 includes RBC, HCT, MCHC, HGB, Iron, MCH and MCV, all related to red blood cell function, as well as AST and ALT, proxies for stress to the liver. This group is anticorrelated with the group of biomarkers in the top left of Fig 9, including ACNeutrophils, WBC, SEGS, Platelets, ACMonocytes and Monocytes, all relating to other cell types.

Other groups can also be seen, such as the group near the center of Fig 9, including AlkPhos, InorgPhos, Lymphs, and ACLymphocytes, relating the phosphate and phosphatase concentrations to lymphocyte count.

#### Age-associated interactions

To search for age-associated interactions, we checked if there were significant differences between the older dolphins (older than 30) and the middle-aged dolphins (between 10 and 30 years old). The results for the type-A interactions are shown in Fig 10 (with the type-B correlations shown in the Supplementary Material). While we saw very few significant differences in the directed interactions between the male- and female-dolphins overall in this data-set (the only exceptions being the influence of magnesium on platelets and inorganic phosphate), several previous studies have found significant differences between the sexes [33, 64, 26, 34]. Consequently, we repeated our analysis after segregating by sex. These results are shown in Figs 11 and 12.

**Figure 10:**
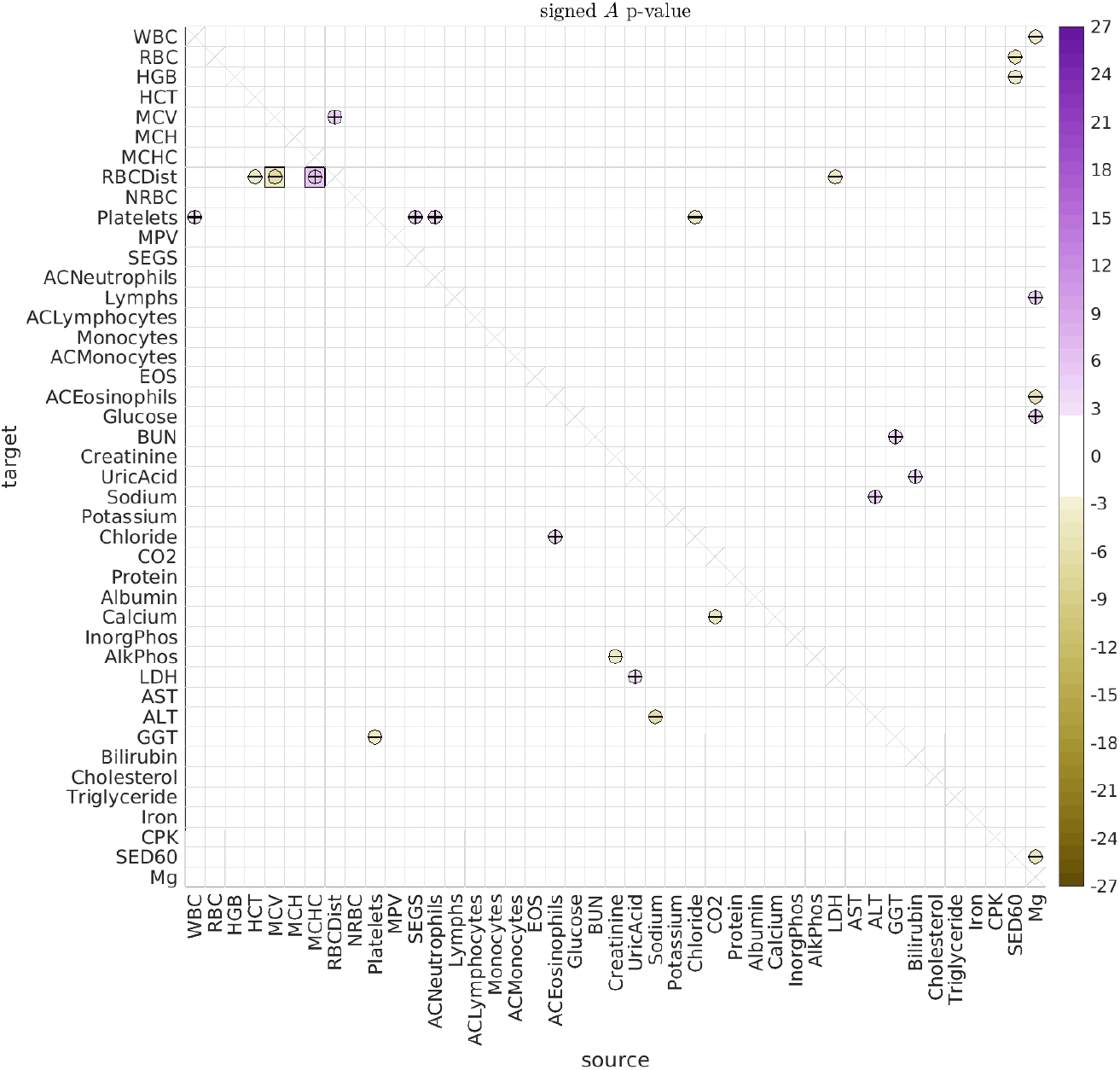
This figure illustrates the significant differences in the directed interactions *A_vv′_* between (i) dolphins over age 30 and (ii) dolphins between the ages of 10 and 30. Positive signs indicate that the corresponding interaction is more positive in the first group (i.e., dolphins over 30) than in the second group (i.e., dolphins between 10 and 30). Negative signs indicate the reverse.

**Figure 11:**
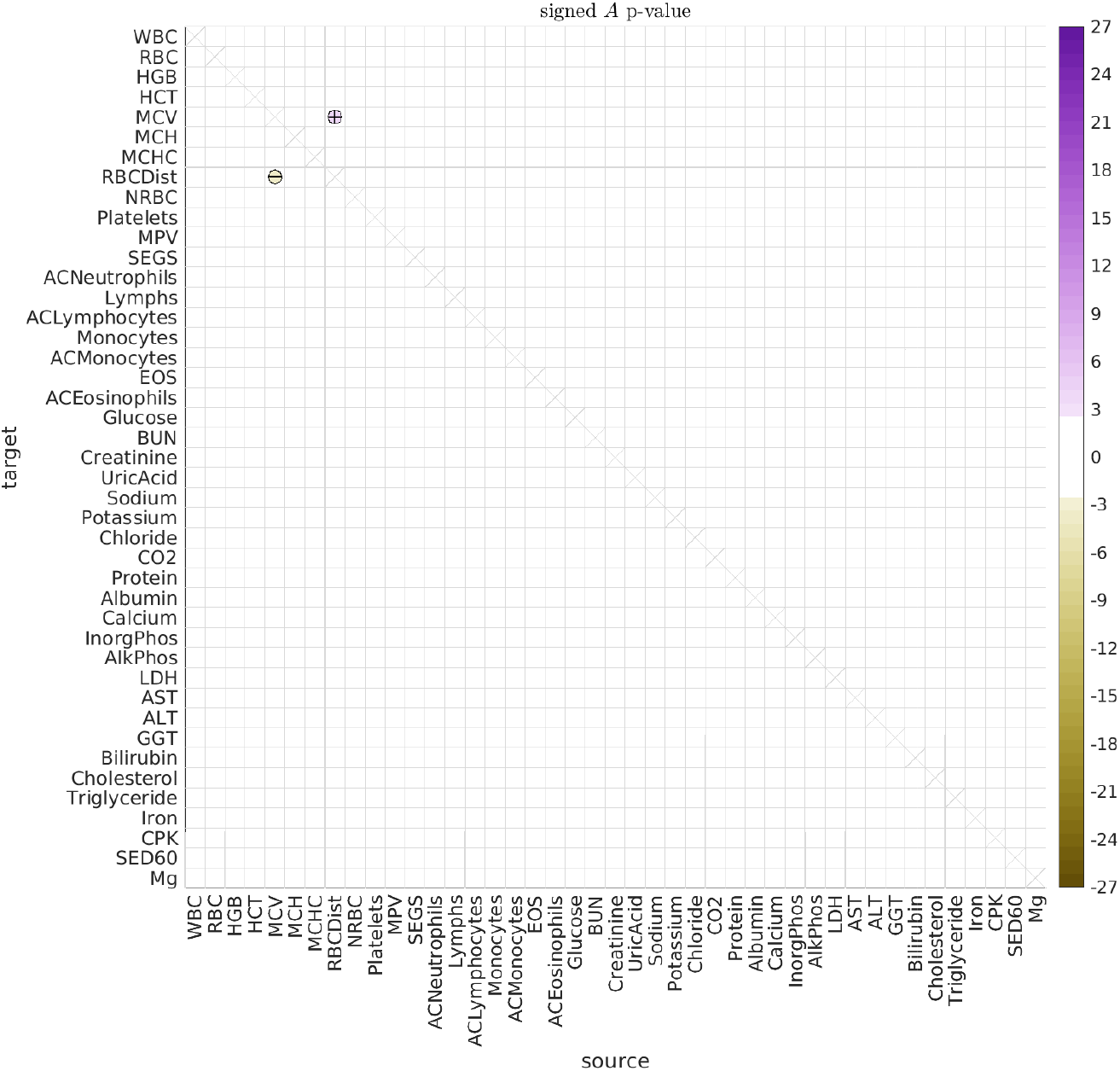
Here we illustrate significant differences in the directed interactions *A_vv′_* between (i) male dolphins over age 30 and (ii) male dolphins between the ages of 10 and 30.

**Figure 12:**
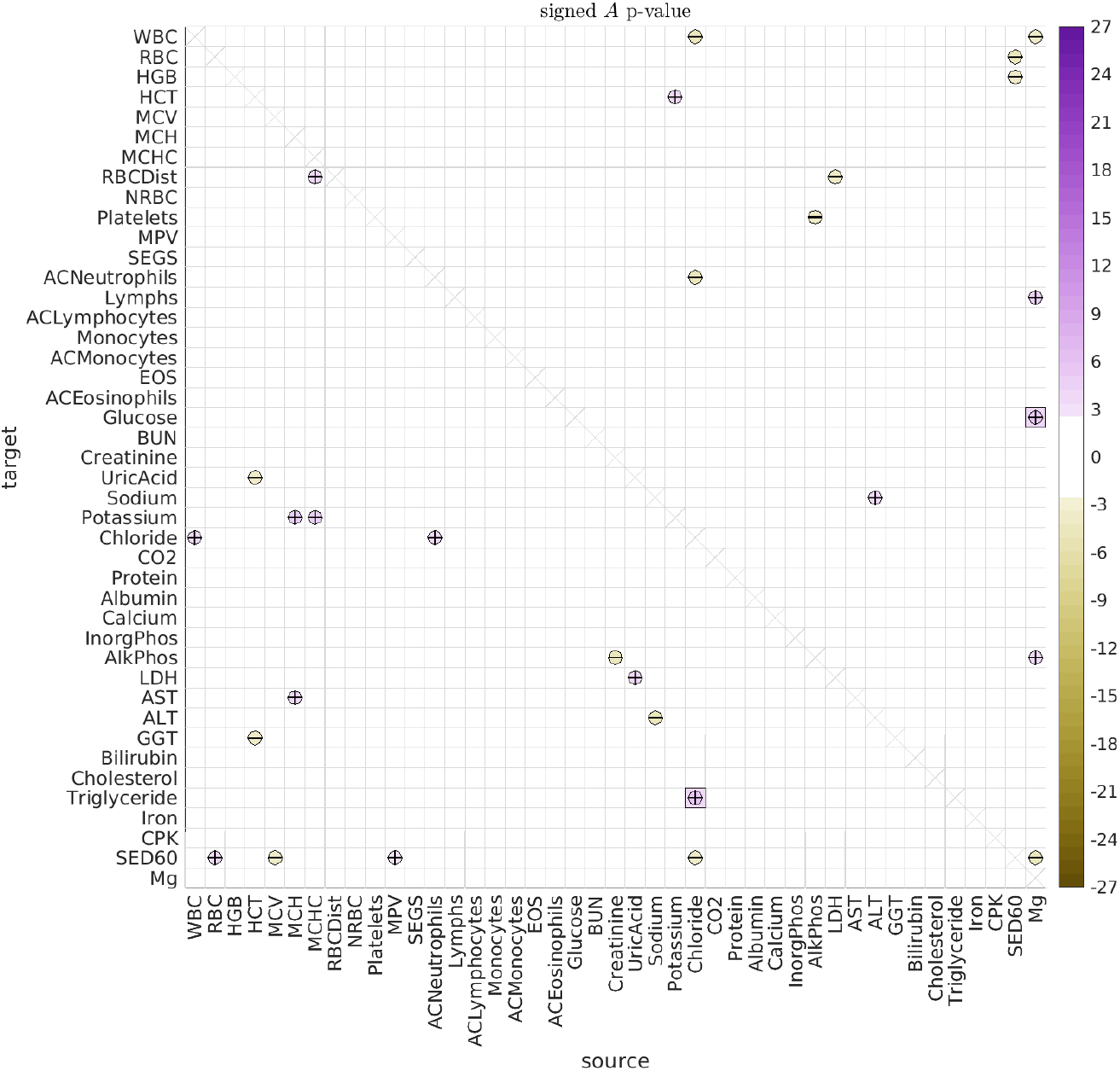
Here we illustrate significant differences in the directed interactions *A_vv′_* between (i) female dolphins over age 30 and (ii) female dolphins between the ages of 10 and 30.

While our sex-segregated analysis has lower power than when the sexes were pooled, we can already see that many of the significant differences observed in the general population are in fact driven by one sex or the other. For example, the male dolphin population drives the age-related change in the push-pull directed interaction between RDW and MCV. On the other hand, the female dolphin population drives many of the other observed age-related changes, such as the interaction between MCHC and RDW, and the interaction between Magnesium and Glucose.

The age-related changes we observe between RDW and MCV could reflect a change in the regulation of bone-marrow function, or the onset of certain kinds of iron-deficiency anemias [65, 66, 67, 68]. More generally, RDW plays a role in various cardiovascular diseases, and it is possible that RDW serves as a proxy for the body’s ability (or lack thereof) to tightly regulate certain processes [69, 70, 71]. Moreover, the management of RDW seems to be important for aging, with at least one study reporting a larger effect-size in men [72, 73].

Regarding the observed interaction between Magnesium and glucose, magnesium is known to affect blood glucose levels and glucose metabolism in humans, and was found to influence glucose absorption in rats [74, 75, 76]. Moreover, magnesium can play the role of a secondary messenger for insulin action, and magnesium accumulation is in turn influenced by insulin [77, 78]. Aging is also associated with changes in magnesium level, and magnesium deficits may result from many age-related conditions [79]. As just one example, the modulation of magnesium levels in elderly patients can potentially improve both insulin response and action, affecting blood glucose levels in turn [80].

## 5 Discussion

We have used a linear SDE to model the time-series data from a longitudinal cohort of dolphins. Importantly, this model accounts for stochastic correlations between biomarkers, without which we would not have been able to accurately estimate the directed interactions.

After performing our analysis, we observed many statistically significant interactions across the population as a whole. Some of the most significant interactions involve ‘push-pull’ biomarker-pairs, where one biomarker excites the other while being inhibited in return. A large block of push-pull interactions involves (i) RBC, HGB and HCT interacting with (ii) AST, MCH, Bilirubin, ALT, Sed60 and Iron, possibly relating to the relationship between liver function and RBC regulation.

We also reported several significant age-related changes to the interaction patterns. Some of the most significant age-related effects involve RDW, MCV and MCHC (in males and females), and the connection between magnesium and glucose (in females).

While certainly suggestive, our a-posteriori analysis cannot be used to decide whether these directed interactions are truly causal or not. Indeed, any significant type-A relationship between any pair of biomarkers (such as Iron and MCH shown in Fig 4) might be due to an unmeasured source affecting both biomarkers, but with different latencies. In this scenario both biomarkers would receive the same unmeasured signal, and would not be causally linked. However, the receiver with the shorter latency would appear to excite the receiver with the longer latency. Moreover, if the unmeasured signal was roughly oscillatory (e.g., meandering back and forth on a time-scale comparable to the difference in latencies), then the receiver with the longer latency would appear to inhibit the receiver with the shorter latency.

In summary, while we believe that most causal interactions would result in a signal captured by our SDE model, the reverse is certainly not true. Causality can only truly be determined after following up our analysis with a controlled study wherein one of the biomarkers is perturbed directly while measuring the response of the other biomarkers.

Finally, we remark that, while dolphins and humans share many age-related qualities [18], they are also different in important ways. For example, female dolphins do not experience menopause, and thus would not be expected to exhibit the suite of hormonal changes that human women do as they age [81]. Further work is needed to determine which of the interactions we have discovered are truly causal, which affect (or are affected by) aging, and which generalize to humans.

## 6 Methods and Materials

### 6.1 Study Population and Animal Welfare

This study was limited to archived, retrospective health data and biospecimen analyses that were collected on Navy bottlenose dolphins (*Tursiops truncatus*) as part of their routine health care. The US Navy Marine Mammal Program is an Association for Assessment and Accreditation of Laboratory Animal Care International-certified program that adheres to animal care and welfare requirements outlined by the Department of Defense and US Navy Bureau of Medicine.

### 6.2 Sample and Data Collection

Archived data for this study were limited to measures obtained from routine blood samples collected from Navy bottlenose dolphins in the morning following an overnight fast between January 1994 and December 2018 (*n* = 5889 samples from 144 dolphins). Blood samples collected either as an initial response or follow-up to acute clinical health concerns were excluded. Methods of routine blood sampling and the measurements obtained from the Navy dolphins have been described previously [18, 64, 35]. Data on the following 44 measures were available for analysis, of which we use *N* = 43: red blood cell indices (RBC count (RBC), hemoglobin (HGB), hematocrit (HCT), mean corpuscular volume (MCV), mean corpuscular hemoglobin (MCH), mean corpuscular hemoglobin concentration (MCHC), RBC distribution width (RBCDist, or RDW), and nucleated RBCs (NRBC)); platelets and mean platelet volume (MPV); white blood cell count (WBC); eosinophils (EOS), lymphocytes (Lymphs), monocytes, and neutrophils (SEGS) (percent and absolute counts, with the latter pre-fixed by ‘AC’); glucose, blood urea nitrogen (BUN), creatinine, uric acid, sodium, potassium, chloride, carbon dioxide (CO2), total protein, albumin, calcium, inorganic phosphate (InorgPhos), alkaline phosphatase (AlkPhos), lactate dehydrogenase (LDH), aspartate aminotransferase (AST), alanine aminotransferase (ALT), gamma-glutamyl transpeptidase (GGT), bilirubin, total cholesterol, triglycerides, iron, creatine kinase (CPK), erythrocyte sedimentation rate (SED60), magnesium (Mg), and estimated glomerular filtration rate (GFR).

### 6.3 Relationships Between Biomarkers

#### Preprocessing

For two of the biomarkers we excluded measurements with atypical lab-codes that lay outside the standard range of measurements seen from the two most common lab-codes. This involved removing Albumin measurements below 2, and RBC distribution width (RDW) measurements above 40. We also ignored the glomerular filtration rate (GFR) biomarker, as for this data-set the GFR measurements *v* are functionally related to the Creatinine measurements *v*′ (up to rounding/discretization error) as:

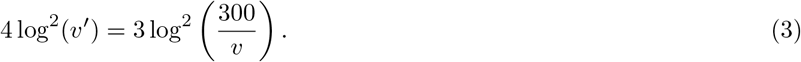

Consequently, after applying the log-transformation described below, these two biomarkers become (almost) exactly correlated with one another.

#### Log-transform

For each biomarker, we applied a log-transformation (adding the smallest discrete increment as a psuedocount) when such a transformation would reduce the skewness of the resulting distribution (across all dolphins). For this data-set the list of biomarkers undergoing a log-transformation is: WBC, MCV, RDW, NRBC, ACNeutrophils, Lymphs, ACLymphocytes, Monocytes, EOS, ACEosinophils, Glucose, BUN, Creatinine, UricAcid, Potassium, Protein, Calcium, AlkPhos, LDH, AST, ALT, GGT, Bilirubin, Cholesterol, Triglyceride, Iron, CPK, SED60, GFR.

#### Normalization

For each dolphin we estimated the age-related drift for each biomarker by using linear-regression on measurements taken from age 5 onwards, excluding outliers above the 99th percentile or below the 1st percentile. We record the slope of this age-related linear drift for each biomarker for each dolphin, referring to this value later on to categorize individual dolphins as slow- or accelerated-agers (see Supplementary Fig 8). After removing this linear drift term, we normalize each biomarker (for each dolphin) to have mean 0 and variance 1.

#### Analysis

For any particular subset of dolphins (e.g., male dolphins between the ages of 10 and 30), we fit a linear stochastic-differential-equation (SDE) to each pair of biomarkers, pooling observations across the dolphins in the chosen subset. Each of these SDEs has 12 parameters in total, accounting for type-A directed interactions, type-B shared stochastic drive, and type-C observation-noise (see (1) and (2)). Briefly, we performed the fit by using a variation of expectation maximization to approximate the model parameters that were most likely to have produced the observed data. The details are described in Supplementary section 3. Our a-posteriori grouping of the type-A directed interactions (shown in Fig 8) is described in Supplementary section 4. The code for these methods can be found within the github repository ‘https://github.com/adirangan/dir_PAD’.

#### Null Hypothesis

To estimate significance for any subset of dolphins across any age-interval for any pair of biomarkers, we compare the estimated parameters of our SDE-model to the parameters obtained after randomly permuting the time-labels of the measurements. Each label-shuffled trial *k* = 1, …, *K* is associated with a permutation of the time-labels *π_k_* which randomly interchanges time-labels within each dolphin (but not across dolphins) within the given age-interval. These label-shuffled trials corresponds to sampling from the null hypothesis that the true interaction coefficient *A*_*vv*′_ for any pair of biomarkers is 0, while still maintaining many of the correlations between biomarkers. Note that the same set of permutations {*π*_1_, …, *π_k_*} is used for the label-shuffled trials 1, …, *K* across all biomarker-pairs *v*, *v*′. Consequently, we can use the correlations between different parameters estimated from the same label-shuffled trial to adjust the p-values of any given parameter (see *p_h_* below).

#### Estimating *p*-values

For each parameter we estimate the *p*-value ‘*p*_0_’ numerically using *K* = 256 label-shuffled trials. When the parameter value for the original data is more extreme than the analogous parameter from any of the *K* label-shuffled trials, we estimate *p*_0_ by first (i) estimating the *z*-score *z*_0_ for that parameter using the Gaussian-distribution fit to the collection of *K* label-shuffled parameter-values, and then (ii) estimating the *p*-value *p*_0_ by applying a 2-sided test to the *z*-score *z*_0_; i.e., 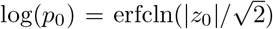. To correct for multiple hypotheses we calculate two adjusted p-values ‘*p_b_*’ and ‘*p_h_*’. The adjusted *p*-value *p_b_* is obtained using a standard bonferroni-correction. Thus, *p_b_* = *p*_0_/*J*, where *J* is the number of parameters under consideration. For the directed interactions *A*_*vv*′_ we set *J* = *N*(*N* – 1), while for the covariances [*BB*^⊤^]_*vv*′_ we set *J* = *N*(*N* – 1)/2 (recall *N* = 43 is the total number of biomarkers analyzed). This bonferroni-corrected *p*-value is an overestimate (i.e., *p_b_* is too conservative). A more accurate *p*-value can be obtained by using an empirical version of the holm-bonferroni adjustment, as described in Supplementary section 5. These strategies can easily be extended to estimate the significance between different groups of dolphins, as described in Supplementary section 5.1.

#### Estimating Aging-rate

To demonstrate consistency with the analysis of [35], we can estimate the aging rate of the dolphins. These results are shown in Supplementary section 6.

## 7 Supporting Information Legends

`SDE_fit_5.pdf:` Description of the methods used in the main manuscript (pdf file).

`Fig_06_A_negative_log_p_bonferroni.csv:` Signed logarithm the bonferroni-corrected *p*-values for A-type interactions shown in Fig 6 (comma-separated values).

`Fig_06_A_negative_log_p_holm_bonferroni.csv:` Signed logarithm the holm-bonferroni-corrected *p*-values for A-type interactions shown in Fig 6 (comma-separated values).

`Fig_07_B_negative_log_p_bonferroni.csv:` Signed logarithm the bonferroni-corrected *p*-values for B-type interactions shown in Fig 7 (comma-separated values).

`Fig_07_B_negative_log_p_holm_bonferroni.csv:` Signed logarithm the holm-bonferroni-corrected *p*-values for B-type interactions shown in Fig 7 (comma-separated values).

`Fig_10_A_negative_log_p_bonferroni.csv:` Signed logarithm the bonferroni-corrected *p*-values for A-type interactions shown in Fig 10 (comma-separated values).

`Fig_10_A_negative_log_p_holm_bonferroni.csv:` Signed logarithm the holm-bonferroni-corrected *p*-values for A-type interactions shown in Fig 10 (comma-separated values).

`Fig_10_B_negative_log_p_bonferroni.csv:` Signed logarithm the bonferroni-corrected *p*-values for B-type interactions associated with Fig 10 (comma-separated values).

`Fig_10_B_negative_log_p_holm_bonferroni.csv:` Signed logarithm the holm-bonferroni-corrected *p*-values for B-type interactions associated with Fig 10 (comma-separated values).

`Fig_11_A_negative_log_p_bonferroni.csv:` Signed logarithm the bonferroni-corrected *p*-values for A-type interactions shown in Fig 11 (comma-separated values).

`Fig_11_A_negative_log_p_holm_bonferroni.csv:` Signed logarithm the holm-bonferroni-corrected *p*-values for A-type interactions shown in Fig 11 (comma-separated values).

`Fig_12_A_negative_log_p_bonferroni.csv:` Signed logarithm the bonferroni-corrected *p*-values for A-type interactions shown in Fig 12 (comma-separated values).

`Fig_12_A_negative_log_p_holm_bonferroni.csv:` Signed logarithm the holm-bonferroni-corrected *p*-values for A-type interactions shown in Fig 12 (comma-separated values).

### 1 Introduction

This is supplemental information for *‘A time-series analysis of blood-based biomarkers within a 25-year longitudinal dolphin cohort’*. In sections 2 and 3 we describe the methods used in the main text to model the longitudinal data. In section 4 we describe the methods used to cluster the array of type-A interactions. These methods are implemented in Matlab within the github repository ‘https://gi-thub.coin/adirangan/dir_PAD’ referenced in the main text. In section 5 we describe our calculation of the holm-bonferroni corrected p-values. in section 6 we describe how we estimate the aging rate of the dolphins.

### 2 Model Structure

As described in the main text, our strategy will be to model the evolution of any *d* specific variables via a simple linear stochastic-differential-equation (SDE). Due to constraints we’ll discuss below, we typically limit ourselves to *d* =2, considering pairs of variables at a time. With this model the evolution over a time-interval [*t*, *t*′] can be approximated as:

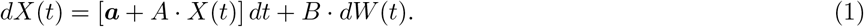

In Eq 1 the vector 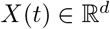 represents the *d*-dimensional vector-valued solution-trajectory at the initial time *t*, the timeincrement *dt* = *t*′ – *t* represents the difference between the initial time *t* and the final time *t*′, and 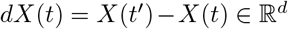 represents the vector of variable-increments between times *t* and *t*′. The vector 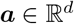 corresponds to a constant ‘velocity’ for each of the variables. The matrix 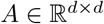 represents the deterministic type-A (linear) interactions between variables. The type-B variation is modeled by the Brownian-increments *dW*(*t*), each drawn independently from the Gaussian distribution 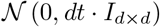. The symmetric matrix 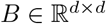 controls the anisotropy of the type-B variation; the covariance of this stochastic term is given by the symmetric-matrix *dtBB*^⊤^ [1]. We assume that we do not measure *X*(*t*) directly, but rather some *Y*(*t*) which depends on *X*(*t*), and which also incorporates the type-C observation-noise:

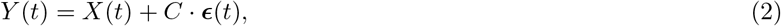

where each vector 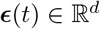 at each observed-time is drawn (independently) from the Gaussian distribution 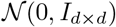. The symmetric matrix 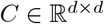 controls the anisotropy of the type-C errors; the covariance of this observation-error is given by the symmetric-matrix *CC*^⊤^.

Note that the observed-times are not necessarily unique: *t*′ could very well equal *t*, and *dt* may equal 0. In this situation *dX*(*t*) ≡ **0**, and so *X*(*t*′) will equal *X*(*t*). However, because of the type-C errors, *Y*(*t*′) will in general be different from *Y*(*t*).

Below we describe the methods we use to fit longitudinal data with the simple dynamical system (SDE) above. After fitting the model parameters we interpret *A* and *BB*^⊤^ as having potential biological significance.

### 3 Implementation

We’ll begin by rewriting Eq 1 slightly. As written above, the velocity ***a*** will contribute a time-dependent term of the form ***a**dt* to the evolution of *X*(*t*). Without loss of generality, we can account for the velocity ***a*** by subtracting a polynomial *Q*(*t*) from *X*(*t*):

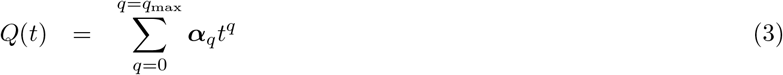

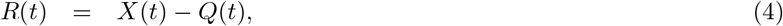

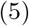

where the 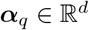 are vector-valued coefficients of the degree *q*_max_ vector-valued polynomial *Q*(*t*). We then assume the term *R*(*t*) evolves according to:

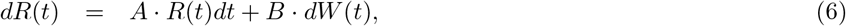

Note that the time-derivative of *Q*(*t*) is given by:

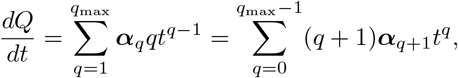

allowing the original representation in Eq 1 to be represented as a special case of Eq 6 simply by setting *q*_max_ = 0 and 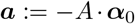. Below we’ll use the term α to refer to the collection of ***α**_q_*; i.e., *α* can be thought of as an array in 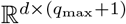.

We’ll also introduce some extra notation to track the different times at which the system is measured, along with their multiplicities. More specifically, we’ll assume that the data itself involves the *j*_max_ observed-times {*τ*_1_, …, *τ*_*j*_max__}, as well as a vector of observations *Y_j_* at each of those times. Because multiple observations might correspond to the same time (e.g., *τ_j_* might equal *τ*_*j*+1_), the number *k*_max_ of unique time-points might be smaller than *j*_max_. To denote these *k*_max_ unique time-points, we’ll use the notation {*t*_1_, …, *t*_*k*_max__}. For example, below we’ll denote the *R*-increments over a time-step via:

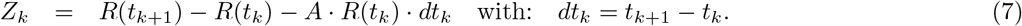

Later on we’ll cross-reference the two arrays for *t* and *τ*, referring to *k*(*j*) as the time-index *k* for which *t_k_* = *τ_j_*.

Before discussing how we fit this model, we remark that we aren’t guaranteed to measure all *d* components of *Y_j_* at each observed-time *τ_j_*; some components of *Y_j_* may be ‘missing’. These missing entries can be treated naturally, and we’ll address this further below in section 3.2.

The various parameters in the model that we’ll consider include *α, A*, *B*, and *C*. The model also involves the hidden (or ‘latent’) trajectory *X*(*t_k_*), corresponding to the ‘true’ (but unknown) system-state at times *t_k_*. The values of *α* and *A* are functionally related to the hidden trajectories *Q*(*t_k_*), *R*(*t_k_*) and *X*(*t_k_*) as described in Eq 6. Finally, the hidden trajectory *X*(*t*_*k*(*j*)_) is related to the observed measurements *Y*(*τ_j_*) via Eq 2.

These quantities can be related within a standard bayesian framework:

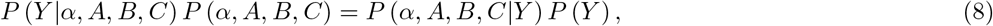

with the likelihood *P*(*α, A, B, C|Y*) defined via:

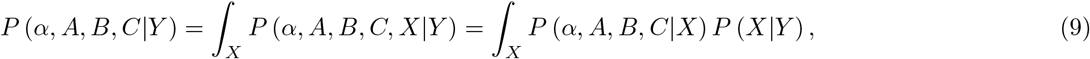

corresponding to an integral over all possible hidden trajectories *X*(*t*).

If we were to assume a uniform prior for the model parameters, then *P*(*α, A, B, C*) would be a constant and we would have the familar expression:

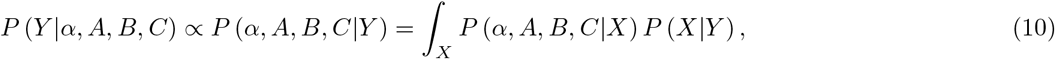

i.e., the probability of the data given the model is proportional to the probability of the model given the data (after marginalizing with respect to the hidden trajectory *X*(*t*)). Thus, to maximize the likelihood of the model we need to find parameters which maximize the likelihood *P*(*α, A, B, C|Y*).

In general, maximizing the likelihood in Eq 10 is challenging, as it is a non-convex function of the model parameters. To approximate a maximum-likelihood solution, we’ll use an iterative-refinement which can be viewed as a version of expectation-maximization [2].

The two main formulae we’ll use in our method are:

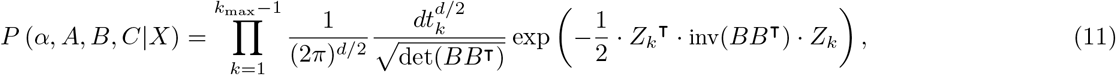

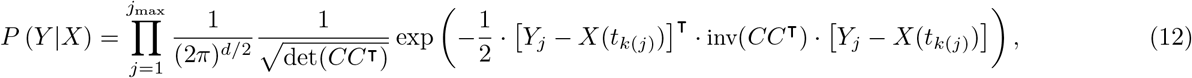

with 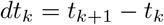 and *Z_k_* defined in Eq 7 and *k*(*j*) corresponding to the *t_k_* that equals *τ_j_*.

By combining Eqs 11 and 12 and using bayes-rule (with a uniform prior on the hidden trajectory *X*), we can approximate the likelihood in Eq 10 via:

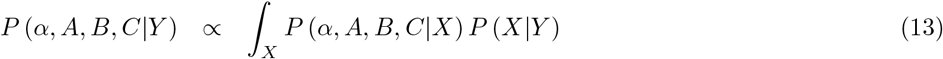

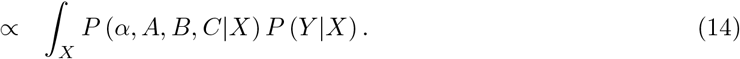

The logarithm of the integrand on the right-hand-side of Eq 14 is a quadratic function of the hidden trajectory *X*(*t_k_*). Consequently, the integral over X in Eq 14 corresponds to a standard Gaussian integral, and can be calculated easily for any fixed values of *α, A, B* and *C*.

Generally speaking, we approximate the model parameters *α, A, B* and *C* iteratively. Referring to the iteration index as *μ*, our algorithm proceeds as follows:

**initialization:** We set *μ* ≔ 0, and initialize the array *α*^[*μ*]^ and matrix *A*^[*μ*]^ to be zero and the matrices *B*^[*μ*]^ and *C*^[*μ*]^ to be the identity.
**update model parameters:** We use nelder-meade optimization to update the model parameters *α*^[*μ*]^, *A*^[*μ*]^, *B*^[*μ*]^ and *C*^[*μ*]^ to maximize the likelihood in Eq 14. Along the way we track the maximum likelihood estimate for the hidden trajectory *X*^[*μ*]^.
**termination:** We stop when the likelihood converges, with a relative-error specified by a global tolerance (e.g., less than 10^−6^).

#### 3.1 Additional Details

Our implementation also includes the following modifications which help to accelerate convergence:

**preliminary optimization:** After initialization, but before beginning nelder-meade optimization, we perform several preliminary rounds of ‘sequential’ optimization. As an example, *C* can be updated by fixing the other model parameters *α*^[*μ*]^, *A*^[*μ*]^ and *B*^[*μ*]^, assuming that *X*^[*μ*]^ is fixed at its (current) maximum-likelihood estimate, and then setting *C*^[*μ*+1]^ to be the value of *C* that maximizes the likelihood in Eq 14. We use this strategy for each of the model parameters, updating *C, B*, *α* and *A* in sequence. We perform this sequential optimization until the likelihood converges (which typically takes 3-4 rounds of updates when *d* =2).
**regularization:** When performing the optimization mentioned above, we subtract regularization terms from the loglikelihood. These regularization terms penalize the magnitude and eccentricity of inv(*BB*^⊤^) and inv(*CC*^⊤^), and can be thought of as non-uniform priors for *B* and *C*. The regularization term 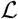 for *B* is:

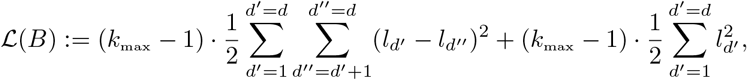

where {*l*_1_, …, *l_d_}* are the *d* log-eigenvalues of inv(*BB*^⊤^). The regularization term for *C* is similar:

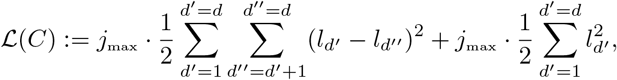

with the *l*-terms now corresponding to log-eigenvalues of inv(*CC*^⊤^).
**regression:** As an alternative to the explicit regularization above, one can estimate *CC*^⊤^ using the empirical covariance of the vectors [*Y_j_* – *X*_*k*(*j*)_]. Similarly, one can estimate *BB*^⊤^ by applying linear regression (with dependent-variable *dt_k_*) to the empirical covariances of the *Z_k_*. In our experience this strategy often produces results which are quite similar to the explicit regularization described above.

#### 3.2 Dealing with missing measurements

As alluded to above, there are observed-times *τ_j_* for which only some of the components of *Y_j_* are observed. When calculating the likelihood for these times we average (i.e., marginalize) over the possible values for the missing entries.

As an example, consider a particular *τ_j_*, corresponding to the time *t*_*k*(*j*)_. The associated likelihood for this term in Eq 12 was originally:

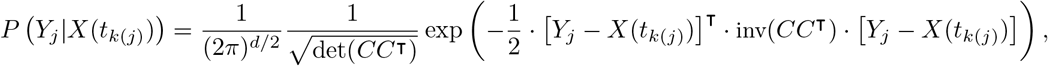

which, after defining 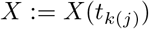 and dropping the index *j* for readability, looks like:

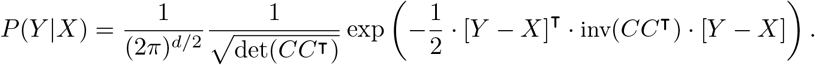

Now let’s assume that only the first *d*′ < *d* components of *Y* are observed. We can write 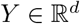 as the concatenation of 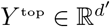 (which is known) and 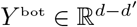 (which is missing):

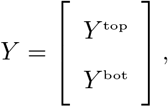

doing the same for *X*. With this notation, the marginalized version of *P*(*Y|X*) for this observed-time becomes:

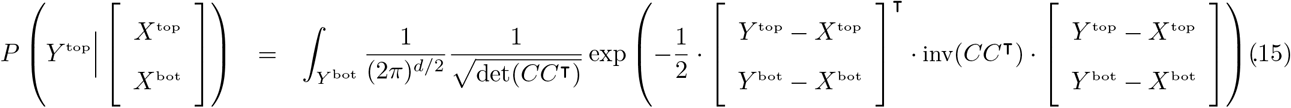

The expression in Eq 15 is a standard Gaussian integral which can be calculated easily when *C* is fixed. When treating multiple missing values for *Y_j_* across different observed-times *τ_j_*, we simply replace each of the corresponding terms in Eq 12 with an appropriately marginalized version analogous to Eq 15.

#### 3.3 Identifiability

Fitting the model above to a particular data-set involves maximizing the likelihood shown in Eq 14. Because this is a nonconvex optimization problem, our strategy is not guaranteed to succeed. Generically, nonconvex optimization of this kind becomes more difficult as the number of model parameters increases. Nevertheless, given the power available within the dolphin data-set, we believe that our implementation can often recover useful information when the number of model parameters is sufficiently low (i.e., when *d* =2 and we are dealing with only pairs of variables).

To demonstrate the effectiveness of our implementation we perform a numerical experiment involving multiple trials. For each trial we fix *d* =2 (corresponding to a pair of variables), set 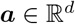 to be the zero-vector, and select 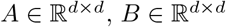 and 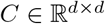 randomly. When randomly selecting *A* we constrain the eigenvalues of *A* to have negative-real-part (so that the resulting SDE produces bounded trajectories) and fix the frobenious norm of *A* to be 1. When randomly selecting *B* and *C* we enforce symmetry, but do not constrain the magnitude of *B* and *C* (i.e., *B* and *C* can have frobenious norm less than or greater than 1). After selecting *A*, we generate a random trajectory *X*(*t*) from the SDE shown in Eq 1 (i.e., using a randomly chosen realization of the weiner process *W*(*t*)). We then sample observed-times from this trajectory, using both the number and distribution of observed-times from the dolphin data-set. Thus, we sample ~ 5300 observed-times *τ_j_*, corresponding to ~ 5200 distinct times *t_k_*, with nonzero time-steps Δ*t_k_* = *t*_*k*+1_ – *t_k_* distributed roughly exponentially with a mean of 0.257 years (see Fig A). For each observed-time *τ_j_* we use *X*_*k*(*j*)_ and *C* to sample *Y_j_* as shown in Eq 2. We then remove roughly ~ 2% of the observed measurements at random (i.e., components of the various *Y_j_*), treating these as missing (in accordance with the dolphin data-set).

**Fig A.**
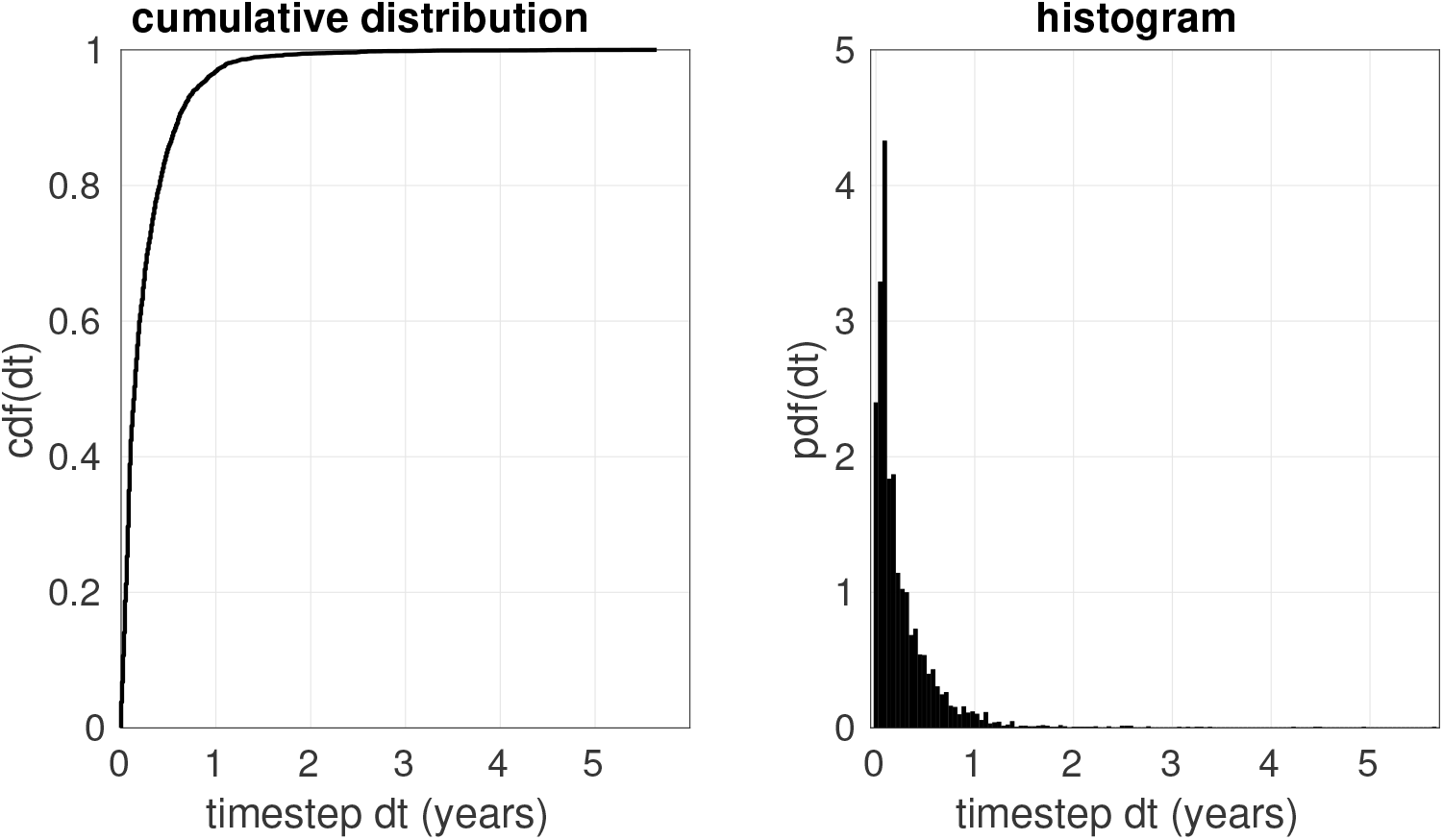
Illustration of the distribution of time-steps within the dolphin data (aggregated across all dolphins). The cumulative-distribution-function is shown on the left and a histogram is shown on the right. This distribution is quite similar to an exponential-distribution with mean ~ 0.257 years. Additionally, for most of the dolphins with many measurements, only a small number (i.e., ~ 2%) of the time-steps are identically zero (corresponding to non-unique observed-times *τ_j_*).

From the data {*τ_j_* } and {*Y_j_*} we then use our methods described above to estimate the model parameters *α*, *A*, *B* and *C* (with *q*_max_ = 0). Once we have estimated the model parameters, we compare the estimated results to the true parameter values used for that trial. For each trial we measure the correlation *ρ*(*A*) between the estimated- and true-values for *A*. We also measure the ratio *σ*(*A*; [*B, C*]) between the frobenius-norm of the true value of *A* and the frobenius-norm of the true value of [*B*, *C*]:

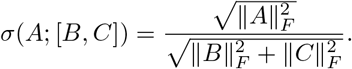

For any particular trial the value of *σ*(*A*; [*B, C*]) can be thought of as a version of a ‘signal-to-noise ratio’, while the value of *ρ*(*A*) measures the recovery quality for *A*.

An example of these results (aggregated over 25600 random trials) is shown in Fig B. For this figure we first bin the trials by their signal-to-noise *σ*(*A*; [*B, C*]), shown along the horizontal. For each of these bins we construct a histogram with respect to *ρ*(*A*). Each column of the heatmap in Fig B shows one of these histograms; the color indicates log2-density (see colorbar on the right). The median of each histogram is indicated in thick cyan, with the 85%-ile and 15%-ile shown in thin cyan. Note that when *A* is roughly the same size as [*B, C*] (i.e., when snr ~ 1, or – log_10_(snr) ~ +0.0) then the recovery is quite good (i.e., close to 100%). When *A* is only one-tenth the size of [*B, C*] (i.e., when snr ~ 1/10, or – log_10_(snr) ~ +1.0) the typical correlation drops to ~ 85% or so. When *A* is only one-hundredth the size of [*B, C*] (i.e., when snr ~ 1/100, or – log_10_(snr) ~ +2.0) the recovery is quite poor, and many trials have a correlation of less than 50%.

We represent the same numerical experiments in a different format within Fig C. In this figure we first divide the trials into three categories. The first category (i.e., ‘*B* small’) corresponds to *B* less than twice the size of *A* (i.e,. ||*B*||_*F*_ ≤ 2). The second category (i.e., ‘*B* medium’) corresponds to *B* between two and eight times the size of *A* (i.e,. 2 ≤ ||*B*||_*F*_ ≤ 8). The third category (i.e., ‘*B* large’) corresponds to *B* more than eight times the size of *A* (i.e,. 8 ≤ ||*B*||_*F*_). These three categories are shown in the left, middle and right subplots, respectively. For each category we measure the frobenius-norm ||*C*||_*F*_, which can be thought of as the inverse of the signal-to-noise ratio relating *A* to *C* (recall that ||*A*||_*F*_ was fixed at 1). For each value of ||*C*||_*F*_ (shown along the horizontal) we again construct a histogram with respect to *ρ*(*A*). Once again, each column showns the log2-density of the associated histogram of *ρ*(*A*) for that value of ||*C*||_*F*_, with the 15, 50 and 85 percentiles indicated in cyan. Note that the recovery of *A* is typically quite good when *B* and *C* are each only a few times larger than *A*. However, when either *B* or *C* is very large, then the recovery of *A* suffers.

In the case of the dolphin data-set, we believe that *B* and *C* are typically between 1 and 6 times bigger than *A*, meaning that 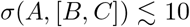, with values of 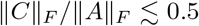. Given the power of the dolphin data-set (i.e., the number of observed-times), this range of model parameters corresponds to typical values of *ρ*(*A*) around 80% or so (see Figs B and C). Thus, we feel reasonably safe reporting our results when *d* = 2.

By contrast, our methods are not sufficiently sensitive to accurately recover the parameters for models involving more than two variables simultaneously. As an example, we repeat the numerical experiments above for the case with *d* = 3 (i.e., three interacting variables). The results are shown in Fig D. Given the power available within the dolphin data-set, we do not typically achieve high recovery quality, even when *B* and *C* are not much bigger than *A*.

### 4 Biclustering the results

In this section we describe the methods we use to identify ‘push-pull’ blocks within the array of type-A interactions. As an example of the structures we are trying to identify, see Fig E.

Our overall strategy is adapted from the ‘loop-counting’ strategy described in [3], giving rise to an unsupervised ‘top-down’ method, and a supervised (or user-informed) ‘bottom-up’ method. The essential idea is to (i) develop a ‘score’ for each row of the array which is correlated with the likelihood that that particular row participates in the push-pull block, and (ii) do the same for the columns.

The top-down method involves calculating these scores across the entire array, then pruning the array by iteratively eliminating rows and columns with the lowest scores. By contrast, the bottom-up method works in the reverse direction, starting with a small collection of user-provided rows and columns (i.e., an initial estimate or ‘seed’ for the push-pull block) and then growing the push-pull block by iteratively adding the rows and columns with the highest scores.

**Fig B.**
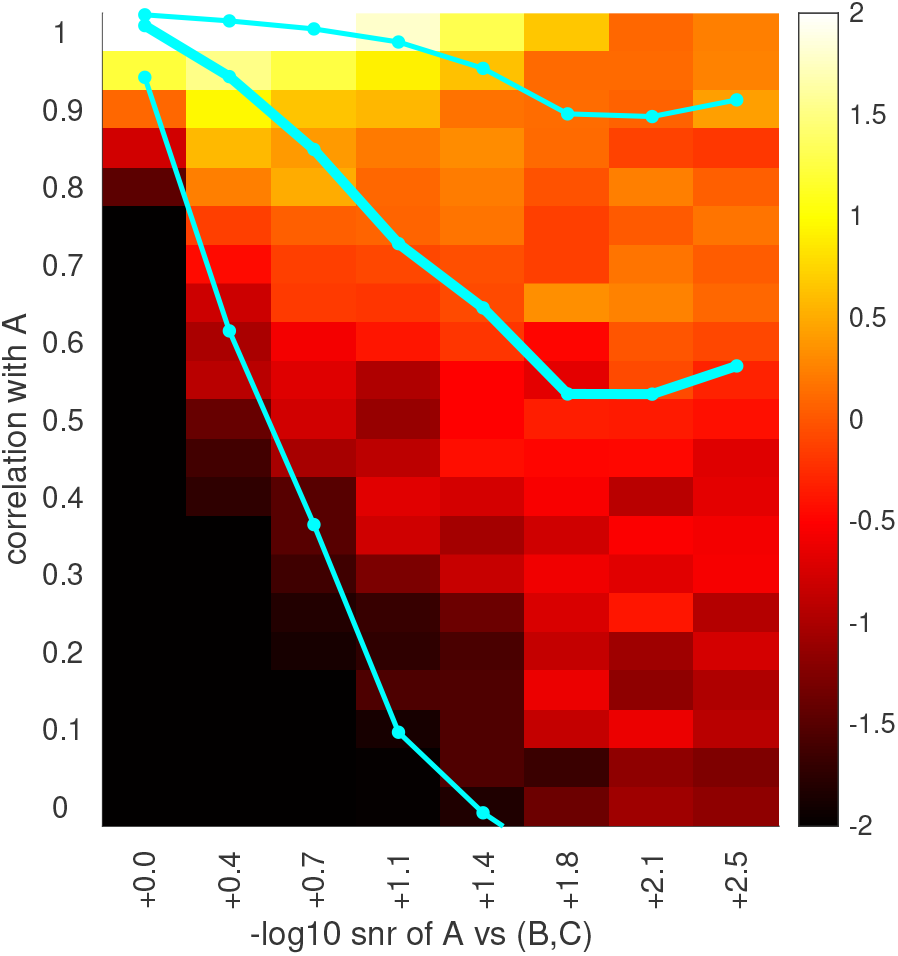
Here we summarize the results of the numerical experiments described in section 3.3 for *d* = 2. The horizontal axis shows a signal-to-noise (snr) comparing the magnitude of the true *A* to that of the true *B* and *C*. For each horizontal location we show a vertical column indicating the histogram of correlations betweeen the estimated and true *A*. The heatmap corresponds to the log2-density of these histograms. The cyan lines indicate the 15, 50 and 85 percentiles for these histograms. Note that when *A* is roughly the same size as [*B, C*] then the recovery is quite good (i.e., close to 100%). However, if *A* is many times smaller than [*B, C*] then the recovery suffers.

**Fig C.**
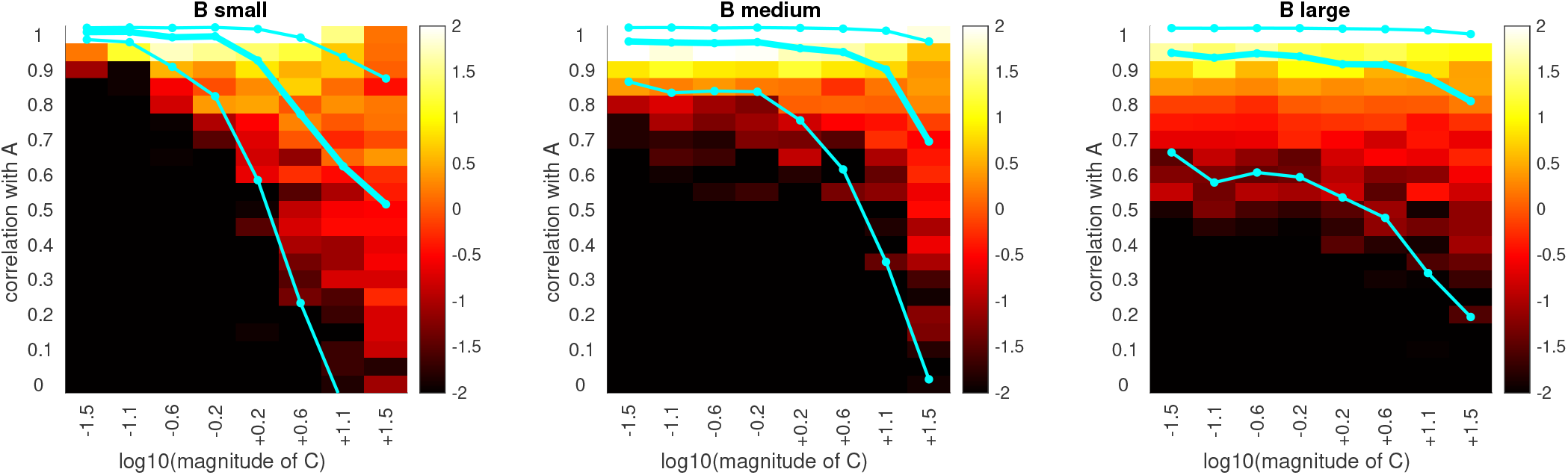
Here we show a different representation of the same numerical experiment from Fig B, involving *d* = 2. This time we divide the trials into three categories, correspond to *B* small, medium and large (left, center and right subplots, respectively). For each category we sort the trials in that category by the frobenius-norm of *C* (shown along the horizontal). For each value of ||*C*||_*F*_ we construct a histogram of the recovery quality (vertical). The 15, 50 and 85 percentiles of these histograms are shown in cyan. Note that the recovery of *A* is typically quite good, except when either *B* or *C* is much larger than *A*.

**Fig D.**
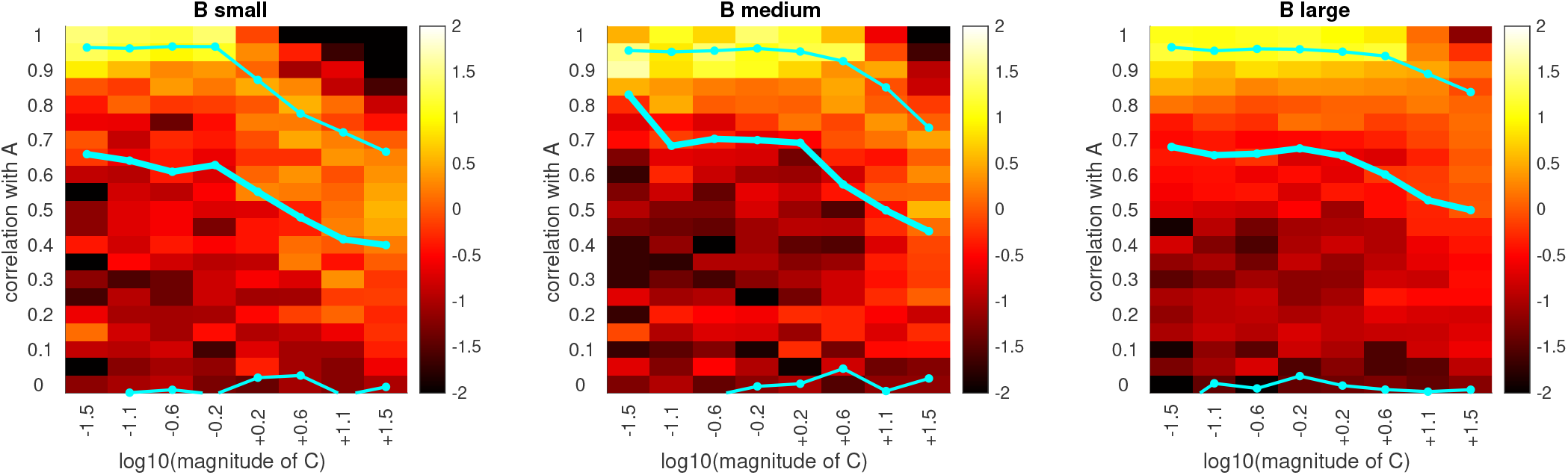
Here we show results for numerical experiments with *d* = 3. The format for this figure is analogous to Fig C. Note that the recovery of *A* is not particularly good, even when both *B* and *C* are of moderate size.

**Fig E.**
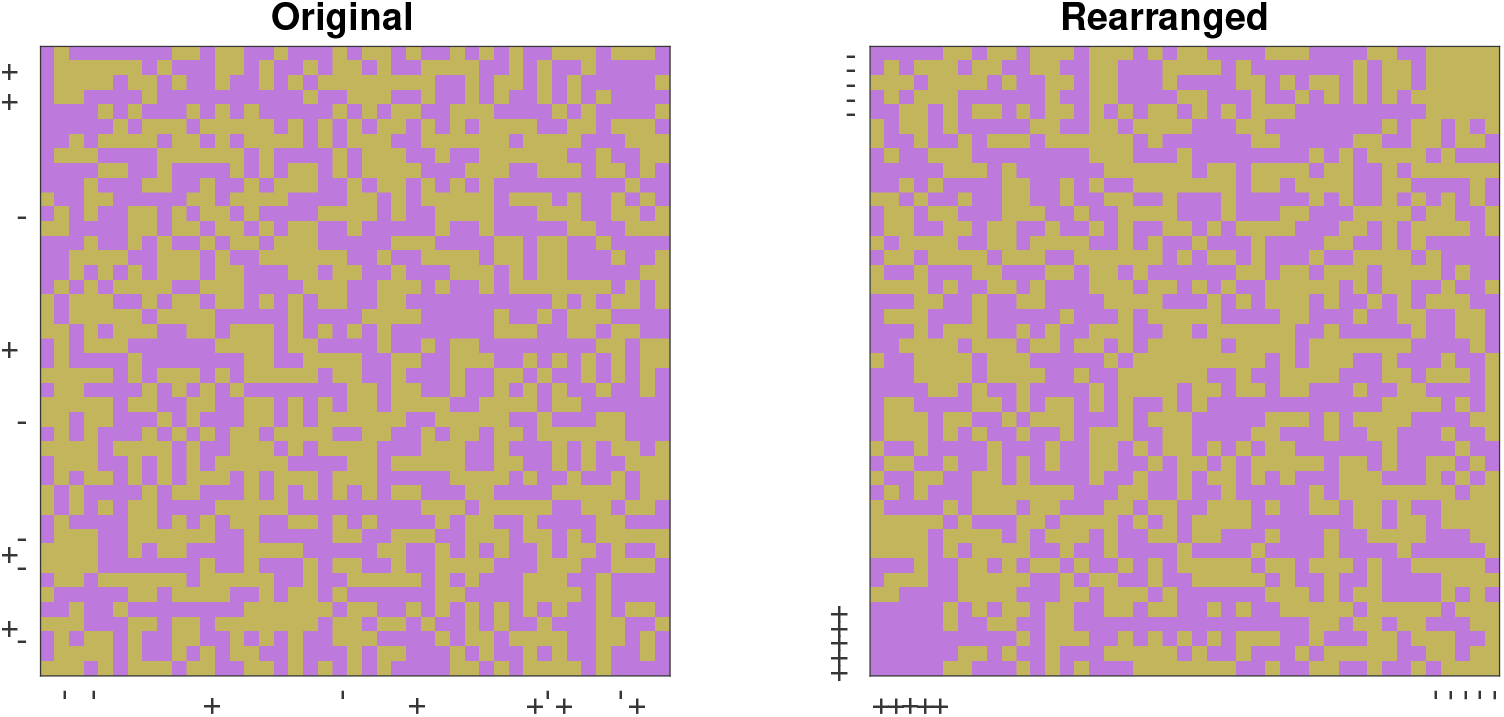
On the right we show a simulated array of type-A interactions. This array is generated by first taking a random matrix (with each entry drawn independently), and then planting a small push-pull block. To define this push-pull block, we first randomly select 2 subsets of 5 variables each, denoted 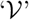 and 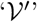, respectively. Once we have defined 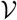 and 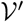, we fix the interactions between any source variable from set 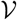 and any target variable from set 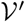 to be excitatory, while fixing the reciprocal interaction to be inhibitory. The variables corresponding to 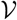 and 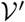 are indicated with the ‘+’ and ‘-’ tick-marks along the axes. If these variables can be identified, then the original array can be re-organized to reveal 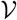 and 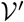 as a pair of contiguous submatrices (see left subplot). In this particular example the original size of the array is *N* × *N*, with *N* = 43, similar to the number of variables used in the dolphin data. The number of interactions within the planted push-pull block is roughly *N*^*M*^, with *M* = 0.5, corresponding to the detection-threshold of our top-down algorithm in the large *N* limit (see [3] and Fig F). We expect our top-down algorithm to reliably find push-pull blocks that are bigger than this threshold when *N* is sufficiently large. We expect our bottom-up algorithm to reliably complete push-pull blocks of this size for a wide range of *N*, assuming that the initial estimate for the push-pull block is a sufficiently large subset of the full block.

In terms of details, we will aim to find sets 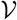 and 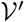 such that the quality 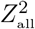 is large, with 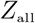 given by:

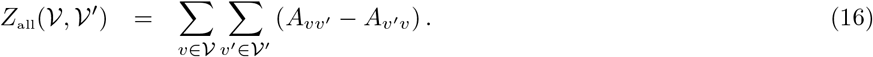

The quality 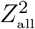 will be large when 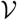 and 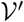 form a push-pull block, with interactions 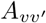 typically of opposite sign to their reciprocal interactions *A_v′v_*.

To search for push-pull blocks, we’ll separate this measure of quality into row-scores:

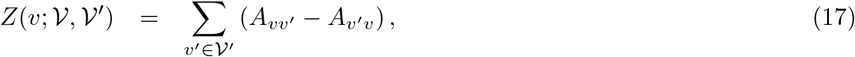

and column-scores:

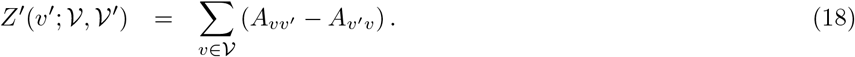

With these definitions one can immediately see that:

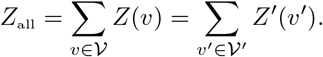

#### 4.1 Top down

These definitions motive a very simple ‘top-down’ method for finding push-pull blocks:

**Initialize:** Define both 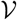 and 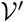 to be the entire set of variables, and set the iteration *μ* = 0.
**Calculate scores:** Use Eq 17 to define row-scores for each variable in 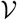 and use Eq 18 to define column-scores for each variable in 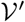. Along the way record the quality 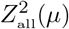 for the current iteration *μ* (i.e., for the current sets 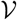 and 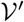).
**Eliminate the lowest scoring variable:** Select one of the elements in either 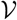 or 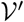 with the lowest score, and eliminate it.
**Iterate:** Iterate this process, recalculating the scores and eliminating the variable (from either 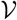 or 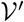) corresponding to the lowest score.

This process will result in a list of variables in the order they were eliminated.

Under certain assumptions, the variables that are retained the longest by the top-down algorithm will form a push-pull block. Because of the similarities between this strategy and the loop-counting methods described in [3], many of the same analytical arguments can be slightly modified to apply to this scenario. For example, the sensitivity of this algorithm is quite similar to spectral clustering [4], and we can make the following statistical claim: If we are given a large random array with a sufficiently large push-pull block hidden within it, then this algorithm will often retain the variables within the push-pull block, eliminating the other variables first. In the limit as the number of variables *N* goes to infinity, this algorithm will succeed with a probability exponentially close to 1 when the number of variables in the push-pull block is 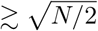 (i.e., when the number of push-pull interactions in the block is ≳ *N*).

**Fig F.**
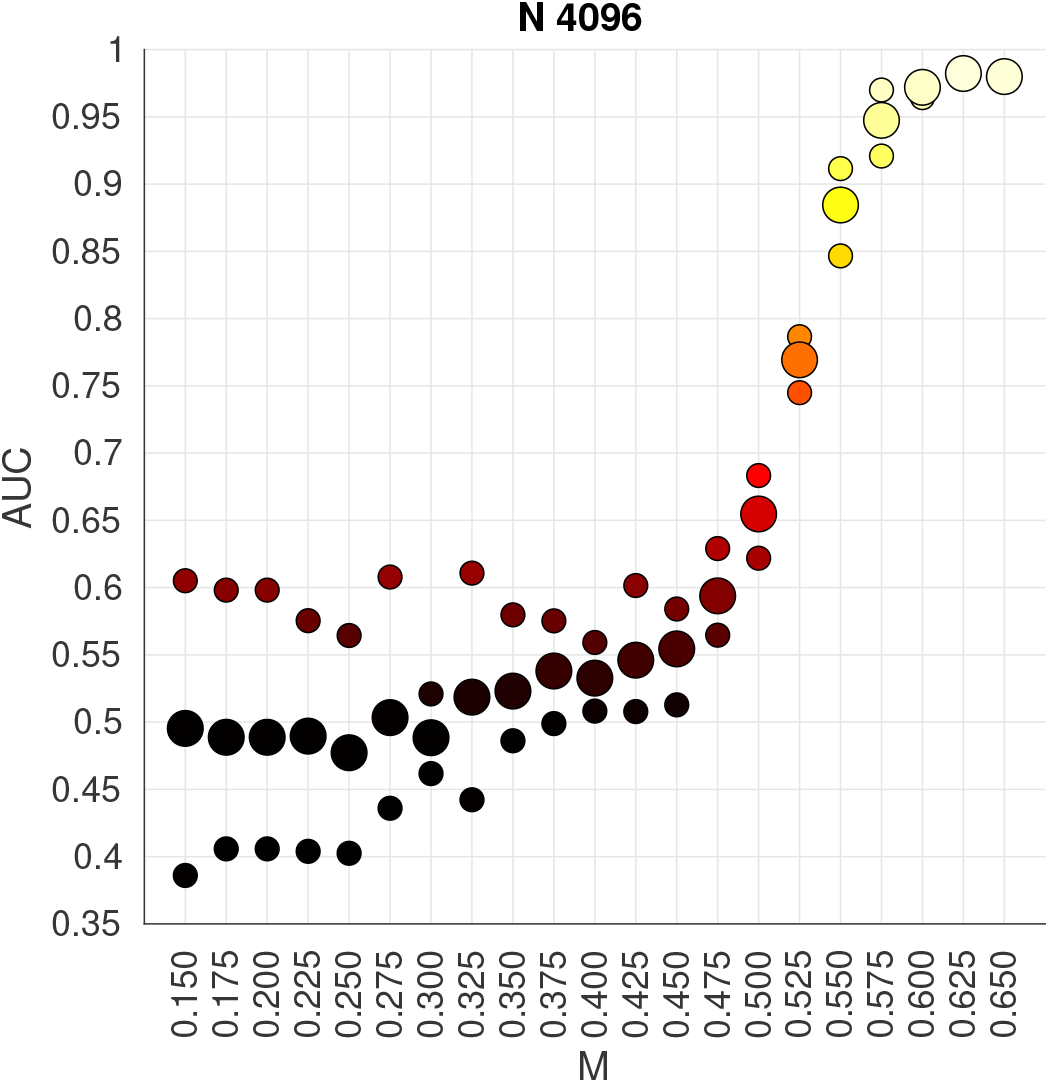
In this figure we illustrate the sensitivity of our top-down loop-counting algorithm for detecting push-pull blocks (see text). For each value of *M* we collect multiple random trials (with different samples of *A*, 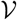 and 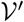) and run our top-down algorithm, measuring the average AUC. The median value is shown with large dots, while the 15%-ile and 85%-ile are shown in smaller dots above and below. Note that the detection-threshold for our algorithm is *M* ~ 0.5, corresponding to push-pull blocks with roughly 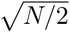 variables within each of them.

To demonstrate this detection-threshold we conduct a numerical experiment. For each trial of this numerical experiment we randomly generate a binary array *A* of size *N* × *N*, with each entry chosen independently from [-1, +1]. Within this array we implant a push-pull block with size 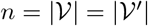 determined by the parameter *M* ∈ [0,1]. For a particular value of *M*, we set 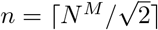, such that 2*n*^2^ ≈ *N*^2*M*^. Thus, the number of push-pull interactions in the planted block is approximately equal to the *M*^th^ power of the total number of interactions in the original array.

Once we have a random binary matrix *A* with a planted block, we run the algorithm described above. We record the list of variables as they are eliminated. After recording this list, we measure the AUC between (i) the listed rank of the variables *not* in 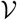, and (ii) the listed rank of the variables *in* 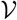. We then measure the AUC’ similarly for 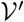, and take the average of AUC and AUC’. If this average AUC is 1, then our algorithm was perfectly successful (i.e., all the variables within the planted block were retained until the very end). If this average AUC is 0.5, then our algorithm performed no better than chance (i.e., the variables within the planted block were not retained any longer than the other variables were).

Results of this numerical experiment are shown in Fig F. This figure plots the average AUC mentioned above as a function of *M*, for *N* = 4096. Note that as *M* approaches 0.5 (and n approaches 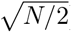), the recovery of our algorithm increases.

In practice, we can determine which variables might form a push-pull block by using a permutation-test. For this permutation-test we compare the quality 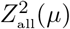 to the distribution of 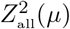 observed when running the algorithm on randomly reorganized versions of the original matrix. The iteration *μ* for which the original quality 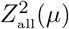 is most significant (relative to the distribution of 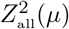 from the randomly reorganized samples) is a natural candidate for the *μ* one should use to identify the entries in the push-pull block. See, e.g., Figs 10 and 11 and the Supplementary Information in [3] for a more detailed description.

#### 4.2 Bottom up

In addition to the top-down algorithm described above, we can also use the row- and column-scores to construct a ‘bottom-up’ algorithm using a strategy similar to many modularity-maximization algorithms (such as louvain clustering [5]):

**Initialize:** Define 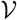 and 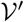 to each be a user-specified group of variables, and set the iteration *μ* = 0.
**Calculate scores:** Use Eq 17 and 18 to define row- and column-scores for each variable in the full set of variables. Along the way record the quality 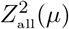 for the current iteration *μ* (i.e., for the current sets 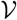 and 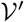).
**Add the highest scoring variable:** Select one of the values of *Z*(*v*) or *Z*′(*v*′) with the highest score, and add the corresponding variable to 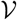 or 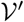, respectively.
**Iterate:** Iterate this process, recalculating the scores and adding variables (to either 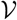 or 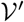) corresponding to the highest score.

This process will result in a list of variables in the order they were added.

Under certain assumptions, the variables that are added first by the bottom-up algorithm will form a push-pull block. For example, if (i) we are given a large random array with a push-pull block hidden within it, and (ii) the inital push-pull block is a sufficiently large subset of the planted block, then (iii) this algorithm will add the variables within the push-pull block first, ignoring the other variables until later. In the limit as the number of variables *N* goes to infinity, this algorithm will succeed with a probability exponentially close to 1 when (i) the initial block is a subset of the planted block, and (ii) the number of variables in the initial block is 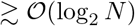.

In practice, we can once again use a permutation-test to determine which variables are within the push-pull block. For this permutation-test we compare the quality 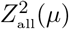 to the distribution of 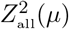 observed when running the algorithm on randomly reorganized versions of the original matrix (where all the entries other than those in the initial block are permuted).

#### 4.3 Analyzing the type-A interactions

We can use the strategies described above to conduct an exploratory analysis of the push-pull blocks within the type-A interactions from the main manuscript. To do so, we first run the top-down algorithm from section 4.1, identifying those variables retained the longest and grouping them into a putative push-pull block. Then we run the bottom-up algorithm from section 4.2 multiple times, using as initial seeds each pair of variables within the putative push-pull block identified earlier. We also run the bottom-up algorithm using as initial seeds each pair of variables that could form a push-pull block themselves (similar in spirit to market basket analysis [6]).

After performing this analysis we recover a large number of slightly different push-pull blocks. Each of these blocks is statistically significant in its own right, but they are far from distinct (indeed, many of these push-pull blocks overlap strongly with one another). While these different push-pull blocks can certainly be merged (using, e.g., the criteria from [7, 8]), we find that the final results of merging push-pull blocks is not robust (i.e., the identity of the final push-pull blocks can change substantially if the merging criteria are altered slightly).

To dodge this issue, we simply step through the list of significant push-pull-blocks, grabbing the largest remaining push-pull block and removing it until nothing of significance remains. This process allows us to order the variables from the original data-set so that several of the most significant push-pull blocks are visually obvious. One such arrangement is shown in Fig 8 in the main text.

When assessing the array of type-A interactions, we can summarize the significance of any particular push-pull block independently, without relying on the methodology used to detect that push-pull block. Given a push-pull block, we define the *p*-value *p_r_* to be the probability that a push-pull block of at least the same size exists within a random array with entries drawn independently (with replacement) from the original A-array. An upper-bound (i.e., conservative estimate) for *p_r_*, denoted *p_u_*, can be constructed using a simple union bound.

To constuct *p*_u_ we introduce the following notation:

1. The original *N* × *N* array of type-A interactions has a fraction *f*_+_ of positive entries, and a fraction *f*_−_ of negative entries. Note that *f*_+_ and *f*_−_ need not necessarily add up to 1.
2. The push-pull block has sizes 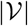 and 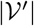 denoted by *V* and *V*′, respectively.
3. The block of interactions {*A*_*v′v*_} between source-variables from 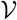 and target-variables from 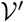 has *n*_+_ positive entries.
4. The block of interactions {*A_vv′_*} between source-variables from 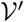 and target-variables from 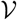 has *n_−_* negative entries.

With these assumptions we can bound *p_r_* via:

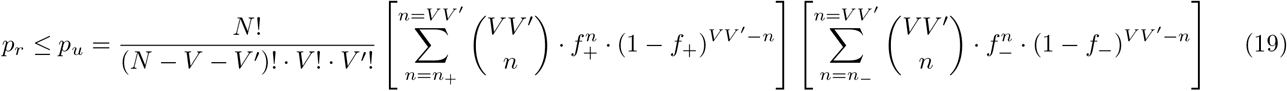

As an example of this upper-bound, we can consider the holm-bonferroni corrected array of type-A interactions for which *f*_+_ and *f*_−_ are each less than 0.10. The push-pull cluster defined by 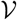 containing RBC, HGB and HCT, and 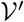 containing AST, MCH, Bilirubin, ALT, Sed60 and Iron has *V* = 3, *V*′ = 6, *n*_+_ = 18 and *n*_−_ = 16; the corresponding upper bound *p_u_* < exp(–50) ~ 2 × 10^−22^. The push-pull cluster defined by 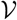 containing AlkPhos and InorgPhos, and 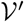 containing CPK, Platelets and BUN has *V* = 2, *V*′ = 3, *n*_+_ = 6 and *n*_−_ = 4; the corresponding upper bound *p_u_* = 0.007. The push-pull cluster defined by 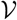 containing Creatinine and Lymphs, and 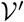 containing AlkPhos, InorgPhos, CPK and Platelets has *V* = 2, *V*′ = 4, *n*_+_ =6 and *n*_−_ = 6; the corresponding upper bound *p_u_* = 0.026. Note that not all visually identifiable push-pull clusters have a low upper-bound *p_u_*. For example, the push-pull cluster defined by 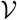 containing MCV, Monocytes and ACMonocytes, and 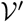 containing RBCDist, Mg and Potassium has *V* = 3, *V*′ = 3, *n*_+_ = 8 and *n*_−_ = 3, yet the corresponding upper bound *p_u_* = 0.28.

### 5 Holm-Bonferroni Adjustment

As mentioned in the main text, the parameters observed for any label-shuffled trial are not uncorrelated with one another. Consequently, the standard bonferroni-corrected *p*-value *p_b_* is an overestimate (i.e., *p_b_* is too conservative). We calculate a more accurate adjusted *p*-value ‘*p_h_*’ by using an empirical version of the holm-bonferroni adjustment.

To describe this in detail consider the collection of *J* = *N*(*N* – 1) parameters *A_vv′_* for all variable-pairs *v* ≠ *v*′. We first determine the *J*-element vector of these parameters for the original-data (by fitting each variable-pair individually). We’ll refer to this vector of parameters as 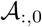, with 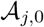 referring to the *j*^th^ parameter from the original data, and the colon ‘:’ referring to ‘all rows’. We then determine the corresponding vector for each of the *K* label-shuffled trials. We’ll refer to each of these vectors as 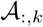 for *k* ∈ 1, …, *K*, one for each trial. Together, the 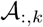 form a *J* × *K* array 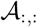, with 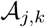 referring to the *j*^th^ parameter from trial *k* (the second colon on 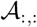 refers to ‘all columns’).

To put these *J* different parameters on equal footing we first convert them all to *z*-scores; for each *j* we convert all the entries of 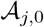 and 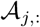 into *z*-scores using the Gaussian-distribution fit to the collection of *K* entries in the row-vector 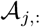. After converting each entry to a *z*-score, we apply a 2-sided test to convert each *z*-score to a negative-log-p-value 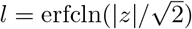. This procedure produces a vector of negative-log-p-values ***l***_*j*,0_ corresponding to the original data, as well as an array ***l**_j,:_* corresponding to the label-shuffled trials. Higher values of *l* indicate more significant values.

Now we sort ***l***_:,0_, as well as each of the ***l***_:,*k*_ in the *j*-direction, in descending order. We’ll refer to these *j*-sorted vectors as 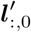 and 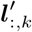.

For each rank-*j* (ranging from the largest and most significant at rank-1 to the smallest and least significant at rank-*J*), we fit the *K* sorted negative-log-p-values 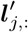 with a gumbel-distribution, denoted by *G_j_*(*l*′). We then use the distribution *G_j_*(·) to ascribe a (one-sided) *p*-value 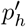 to the value 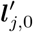. Finally, we set the adjusted p-value *p_h_* to be 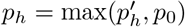.

We can confirm that our holm-bonferroni adjusted p-value *p_h_* is more accurate than the original bonferroni-adjusted *p_b_* by measuring the fraction *P_h_*(*x*) of label-shuffled trials with at least one holm-bonferroni adjusted *p*-value (taken across the *J* variable-pairs) less than *x*. This cumulative-distribution function is quite close to the line *P_h_*(*x*) = *x*, indicating that *p_h_* is an accurate estimate of the true *p*-value, after adjusting for multiple hypotheses (e.g., see Fig G).

#### 5.1 Estimating Significant Differences

To search for significant differences between two different subsets of dolphins, we employ the same strategy described above, except applied to the difference between parameters (rather than the parameters themselves). Additionally, when comparing two different subsets we use all pairs of permutations *π_k_*,*π_k′_* to define the label-shuffled distribution.

To be more explicit, imagine that the first set is ‘dolphins between the ages of 10 and 30’, while the second set is ‘dolphins over the age of 30’. We’ll denote these two sets by *S*^1^ and *S*^2^. Continuing with the example we used above when discussing the holm-bonferroni adjustment, we would measure the *J* parameters *A_vv′_* (across all variable-pairs *v*, *v*′) for each set, denoting the results with the vectors 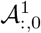 and 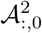, respectively (note that here the superscript does not indicate an exponent, but rather the subset considered). We also do the same for each of the *K* label-shuffled trials, producing the arrays 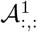 and 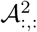. Before proceeding any further we calculate the differences 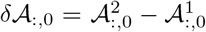 for the original data, and the *K*^2^ differences 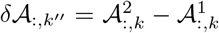, for each pair of label-shuffled trials, where the index *k*″ = 1, …, *K*^2^ enumerates all the trial-pairs *k, k*′.

**Fig G.**
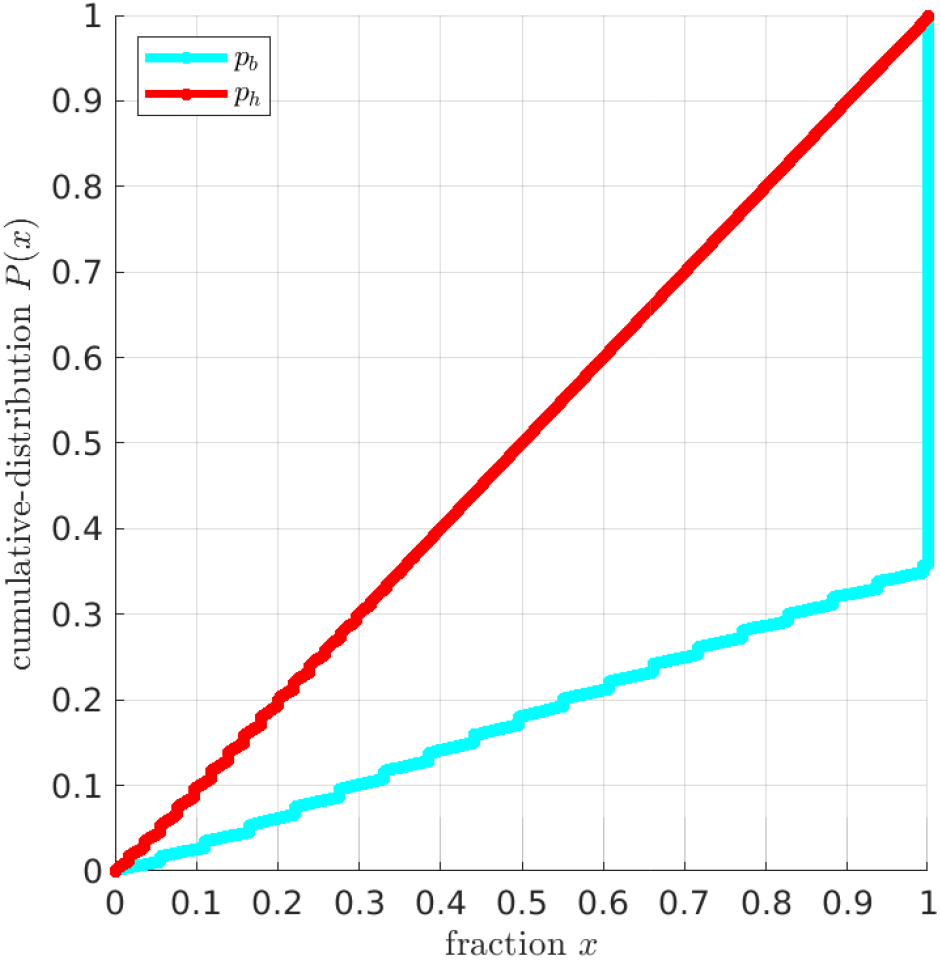
Here we illustrate (in red) the cumulative-distribution *P_h_*(*x*) for the holm-bonferroni adjustment, as calculated for the deterministic interaction terms *A*_*vv*′_ across all variable-pairs. This cumulative-distribution *P_h_*(*x*) is defined to be the fraction of label-shuffled trials exhibiting at least one value of *p_h_* (considered across all variable-pairs) less than *x*. The *p*-values refer to the significance of the difference (in parameters) between the subsets *S*^2^ and *S*^1^. The subset 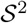 refers to all dolphins over the age of 30, while the subset 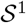 refers to dolphins between the ages of 10 and 30. An analogous cumulative-distribution *P_b_*(*x*) for the bonferroni-correction is shown in cyan. Note that the value of *P_h_*(*x*) closely aligns with the identity line (grey).

From here on we proceed as usual, replacing (respectively) the vector 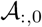 and the *J* × *K* array 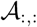 with the vector 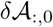 and the *J* × *K*^2^ array 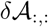. An example of adjusted *p*-values produced using this approach is shown in Fig G.

### 6 Estimating Aging-rate

To demonstrate consistency with the analysis of [9], we measure the slope of the age-related drift for the 6 biomarkers Hemoglobin (HGB), Alkaline Phosphatase, Platelets, Lymphocytes, Creatinine and Protein across each of the dolphins from age 10yr onwards. We project all dolphins onto the first principal-component ‘*u*’ of this array, separating them into two categories based on the median of *u*. Those dolphins with larger-than-median *u*-values typically exhibit slow deterioration of Hemoglobin, Alkaline Phosphatase, Platelets and Lymphocytes, along with a slow accumulation of Creatinine and Protein; these are classified as slow-agers. Conversely, the dolphins with lower-than-median u-values typically exhibit rapid deterioration of Hemoglobin, Alkaline Phosphatase, Platelets and Lymphocytes, along with rapid accumulation of Creatinine and Protein; these are classified as accelerated-agers. As shown in Fig H, the accelerated-agers tend to develop anemia and lymphopenia and clinically low levels of alkaline phosphatase and platelets more rapidly than the slow-agers.

**Fig H.**
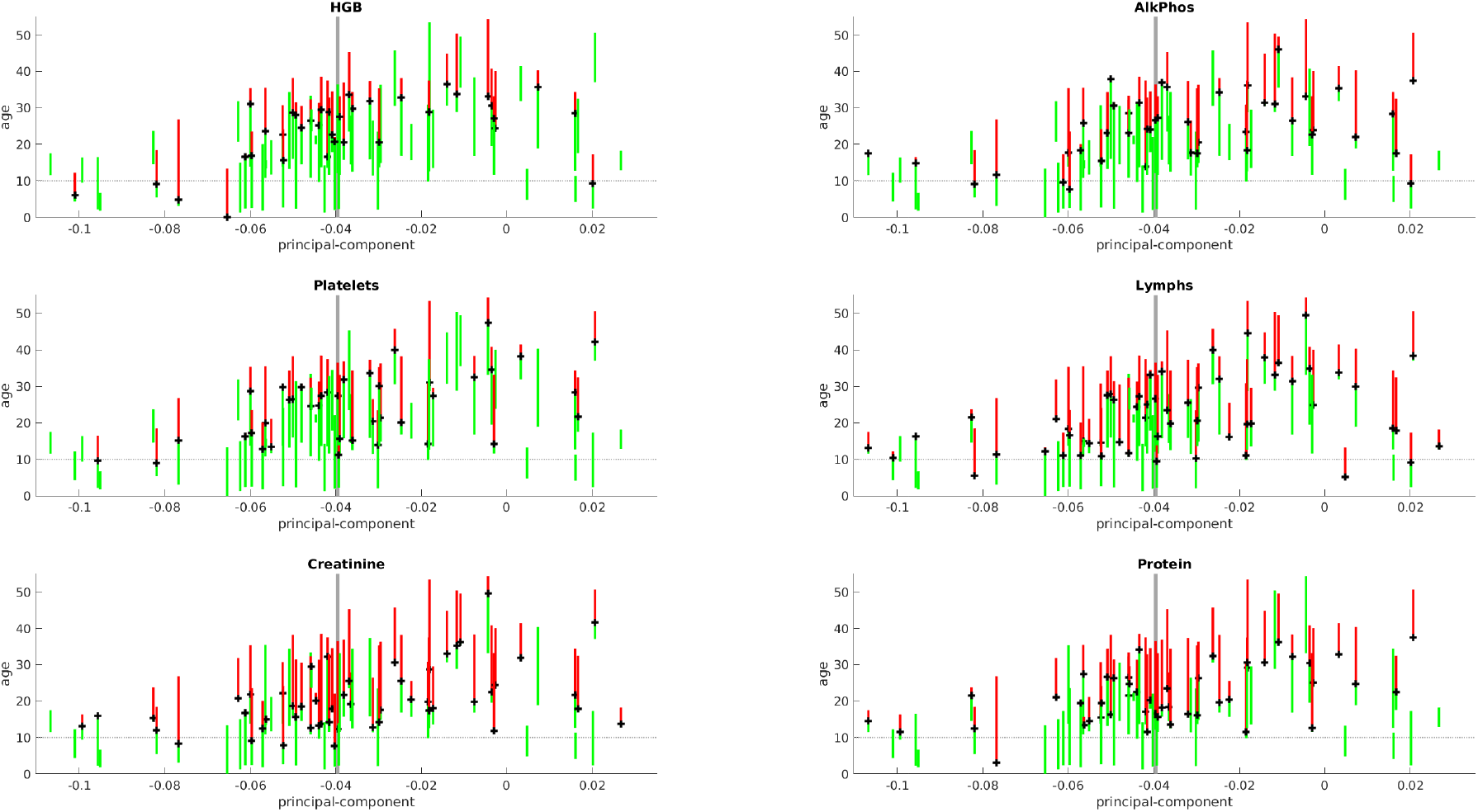
Here we illustrate the distribution of dolphins with regards to aging rate. As described in the Methods, we have measured the slope of the age-related drift for the 6 biomarkers HGB, AlkPhos, Platelets, Lymphocytes, Creatinine and Protein, using only measured ages above 10 years. We calculated the first principal-component *u* of this array, and projected each dolphin onto this principal-component. In each subplot we illustrate the correlation between age-related conditions and the *u*-value for each dolphin. Taking the first subplot (HGB) as an example, each dolphin is illustrated using a vertical line positioned at that dolphin’s u-value. Each vertical line spans the dolphin’s measured ages, with green segments indicating ages where the dolphin exhibits normal values of HGB, and red segments indicating ages after which the dolphin first exhibited a low value of HGB (i.e., anemia). The threshold we use to distinguish normal HGB from anemia is the value of HGB=12 listed in table-2 of [9]. For visual clarity we place a black ‘+’ at the age when each dolphin first exhibits a low HGB value. One can clearly see a correlation between the u-value for each dolphin and the age at which that dolphin first exhibits anemia. The remaining subplots are analogous to the first, referencing the other 5 age-related variables described in table-2 of [9]. In the background of each subplot we highlight the median value of *u* ~ −0.04, which we use as a threshold to categorize dolphins into slow-agers (*u* > −0.04) and accelerated-agers (*u* < −0.04).

**Fig I.**
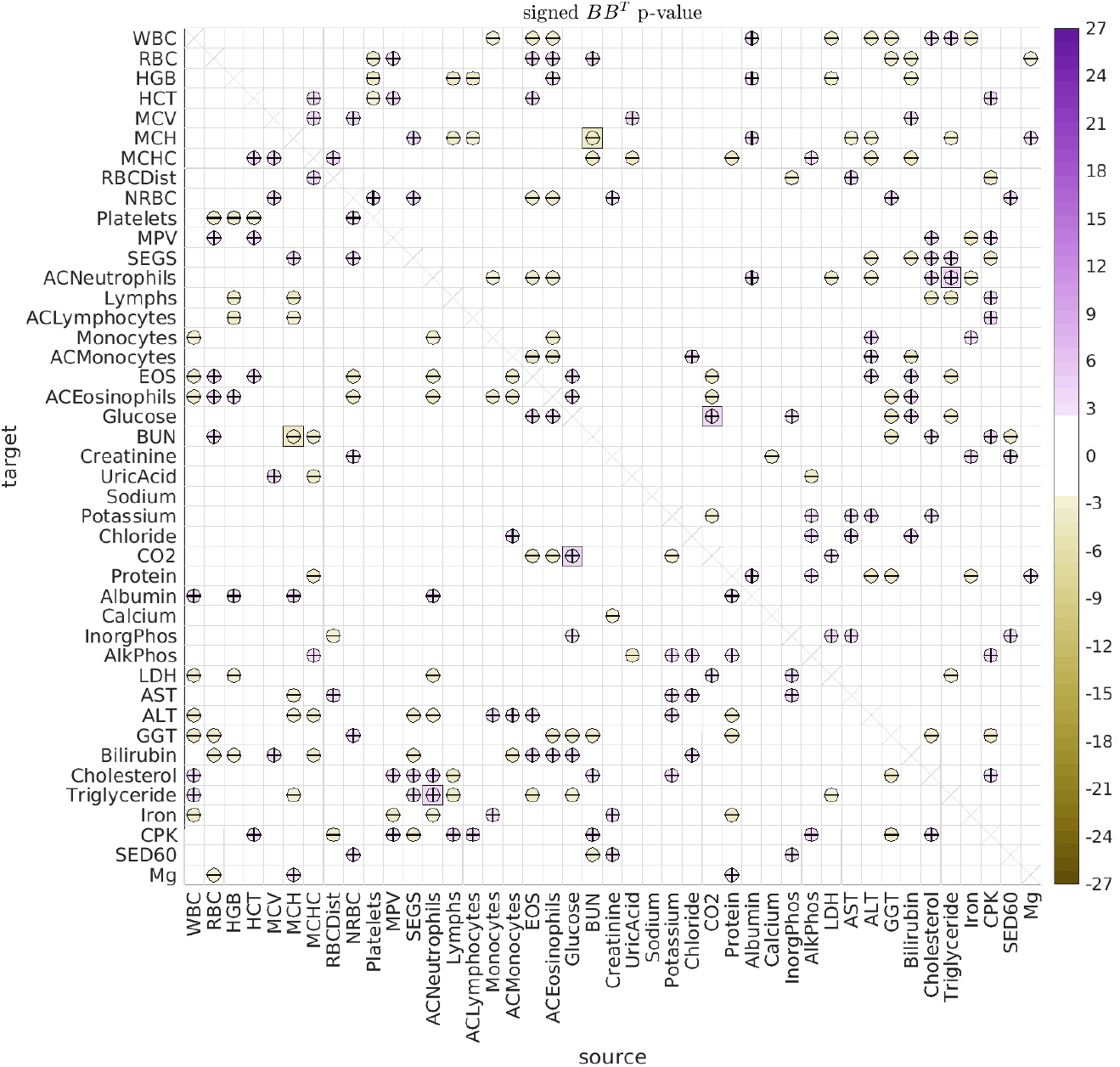
This figure has the same format as Fig 7 in the main text, showing the significant differences in the stochastic correlations [*BB*^⊤^]_*vv*′_ between (i) dolphins over age 30 and (ii) dolphins between the ages of 10 and 30.

**Fig J.**
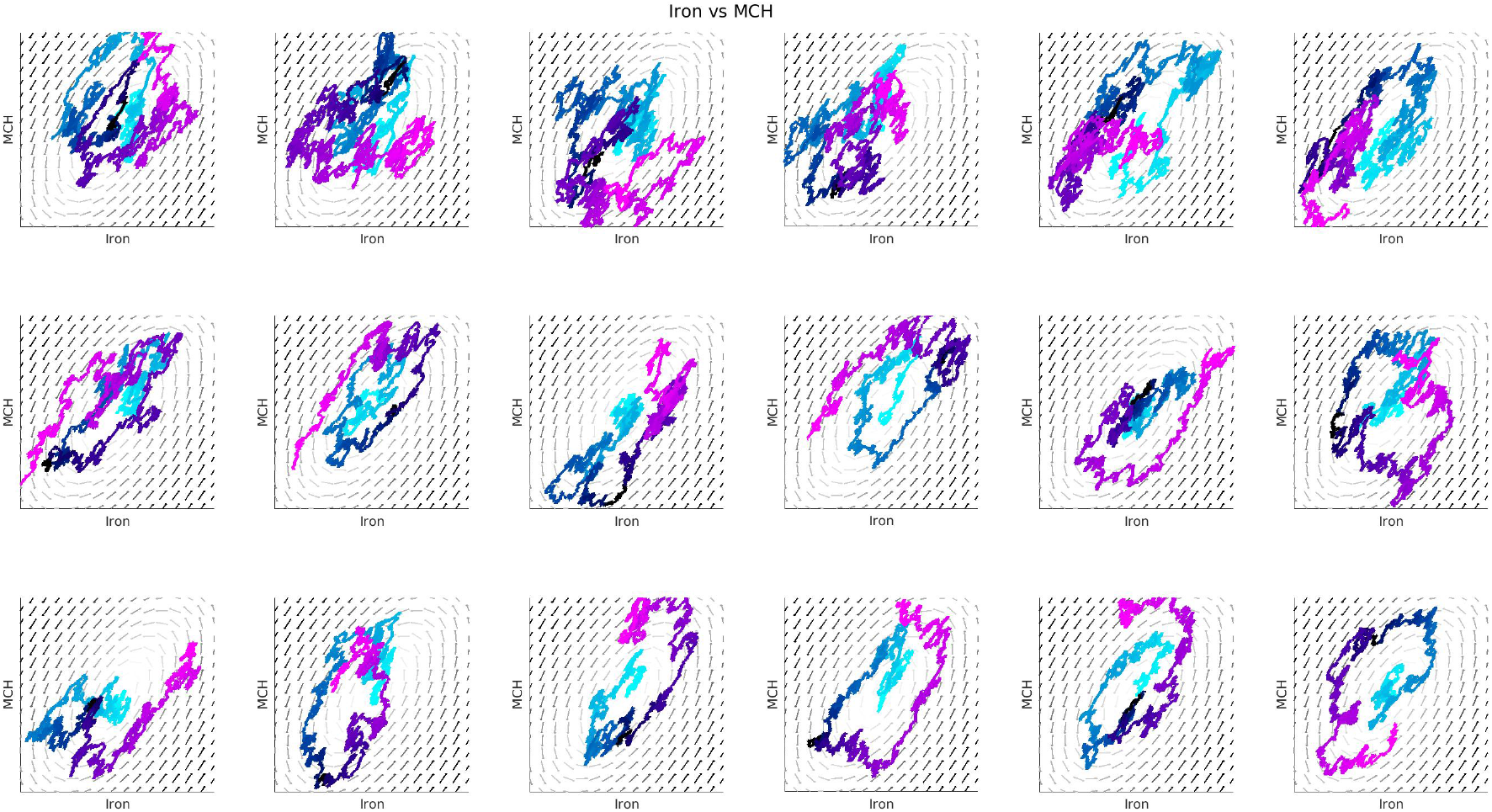
In this figure we show many different realizations of the stochastic differential equation (SDE) relating Iron to MCH. Each panel in this figure has the same format as the left side of Fig 4 from the main text.

